# Kingdom-wide comparative transcriptomics reveals deeply conserved and predictable stress response programs across Viridiplantae

**DOI:** 10.64898/2026.04.17.719161

**Authors:** Eugene Koh, Li Hui Peh, Marek Mutwil

## Abstract

How conserved stress responses are across the plant kingdom remains poorly understood. Here, we present a kingdom-wide stress transcriptome atlas of 36 Viridiplantae species, from chlorophytes to angiosperms, across nine abiotic and biotic stresses. The atlas integrates reanalyzed public RNA-seq data with new in-house stress experiments on three species representing basal lineages, yielding 13.6 million differential expression calls from over 3,200 manually curated control-treatment comparisons. We find that ancient gene families respond broadly but moderately, while lineage-specific families respond narrowly but intensely, revealing a division of labor in stress gene deployment. Stress response conservation decays with phylogenetic distance yet remains detectable across more than 700 million years of divergence, with upregulated genes diverging faster than downregulated genes. Functional co-occurrence analysis uncovers a deeply conserved growth-defence tradeoff alongside stress-specific transcriptional rewiring. Conserved stress co-expression modules undergo regulatory subfunctionalization through duplication, with whole-genome duplicate pairs preferentially retained within modules. Finally, DNA and RNA foundation models predict stress responsiveness from sequence alone (auROC 0.755), suggesting a partially conserved cis-regulatory code underlying stress responses across the kingdom.

## Introduction

Plants cannot relocate to escape unfavourable conditions and must instead rely on transcriptional reprogramming to survive environmental stress. Research over several decades, predominantly in *Arabidopsis thaliana* and a small number of crop species, has identified the core molecular components of plant stress responses: heat shock proteins are induced by high temperatures (Kotak et al., 2007; Lindquist, 1986), the CBF/DREB regulon coordinates cold acclimation (Chinnusamy et al., 2007; Thomashow, 1999), and abscisic acid signalling mediates responses to drought and salinity (Cutler et al., 2010; Zhu, 2002). These findings have shaped a detailed picture of stress gene regulation, but one that is derived almost entirely from fewer than ten species (Koh, Sunil, et al., 2024). A central question therefore remains open: to what extent are stress transcriptional programs conserved across Viridiplantae evolution, and where have they diverged?

Comparative transcriptomics can address this question, but progress has been constrained by taxonomic scope. Cross-species stress studies have generally compared two to five species within a single lineage (Y. Dong et al., 2023; Hartmann et al., 2022) or examined one stress type across a moderate phylogenetic range (Wu et al., 2021). Large-scale expression resources such as the EMBL-EBI Expression Atlas (Moreno et al., 2022), PEO (Koh, Goh, et al., 2024) and CoNekT (Proost & Mutwil, 2018) aggregate data from many species, but were not designed for systematic cross-species comparison of stress responses. The LSTrAP-Cloud pipeline (Tan et al., 2020) and its kingdom-scale extension LSTrAP-Kingdom (Goh & Mutwil, 2021) have made it feasible to process public RNA-seq data uniformly across dozens of species, yet no previous study has applied such a pipeline to construct a curated, kingdom-wide stress atlas with matched control–treatment comparisons. We therefore lack a unified view of how plant stress responses are organized at the gene family level, how they are wired into functional modules, and whether the regulatory logic uncovered in *Arabidopsis thaliana* and other model organisms generalizes across the kingdom.

Three specific gaps limit current understanding. First, it is unknown how gene family age relates to stress response profiles. Ancient, broadly conserved families and recently evolved, lineage-specific families could play overlapping or complementary roles in stress defence, but distinguishing between these possibilities requires expression data across a deep phylogenetic sample. Second, the higher-order functional architecture of stress responses, including which pathways are co-activated, which are co-suppressed, and how this wiring differs between stresses, has not been mapped at the kingdom scale. Studies on growth–defence tradeoffs have documented antagonism between growth and immunity in individual species (Huot et al., 2014; Züst & Agrawal, 2017), but it is unknown whether this tradeoff reflects a deeply conserved transcriptional program or independent evolution. Third, it remains unclear whether stress responsiveness can be predicted from gene sequence alone. DNA and RNA foundation models trained on plant genomes (Yu et al., 2024; Zhai et al., 2025) offer a potential route to predicting stress gene behaviour in species that lack expression data, but this has not been tested across a broad phylogenetic sample.

Here, we address these gaps by constructing a cross-kingdom stress transcriptome atlas spanning 36 species from seven major Archaeplastida lineages and nine stress types. We reanalyzed publicly available RNA-seq datasets using LSTrAP-Cloud with Kallisto for transcript quantification (Bray et al., 2016; Tan et al., 2020), supplemented by in-house stress experiments on *Brachypodium distachyon*, *Selaginella moellendorffii* and *Klebsormidium nitens* to fill gaps in basal lineages. Control–treatment pairs were manually curated for each species and stress, yielding over 3,200 comparisons and 13.6 million differential expression calls within the 36 species.

We first validate the dataset by testing whether canonical stress marker genes respond as expected across species. We then characterize how evolutionary age shapes stress response breadth and intensity, and quantify the decay of transcriptome conservation with phylogenetic distance. At the functional level, we use MapMan-based co-occurrence analysis to identify a conserved growth–defence tradeoff and stress-specific transcriptional signatures, and cross-validate our findings against the PlantConnectome knowledge graph (S. C. Lim et al., 2025). We show that stress co-expression modules are conserved across species and undergo subfunctionalization through duplication, with whole-genome duplicate ohnologs preferentially retained within co-regulated units. Finally, we demonstrate that DNA and RNA foundation models (Yu et al., 2024; Zhai et al., 2025) can predict stress responsiveness from sequence across 25 species, and identify the sequence and structural features that underlie this prediction.

### Kingdom-wide stress transcriptome atlas assembly and validation

Here we have collected and manually curated a stress transcriptome atlas consisting of nine abiotic and biotic stresses. The atlas integrates reanalyzed public RNA-seq data with new in-house stress experiments on three species representing basal lineages, yielding over 3,206 manually curated control–treatment comparisons (Fig. 1A, Table S1). These species were chosen to represent the major clades of Viridiplantae phylogeny, and also the availability of stress experiments present in the Sequence Read Archive database (Leinonen et al., 2011). The dataset coverage is uneven across the phylogeny: model species (*Oryza sativa indica* with 481 pairs, *Arabidopsis thaliana* with 413, *Zea mays* with 248) dominate, while basal lineages (*Selaginella moellendorffii, Klebsormidium nitens*) have fewer than 10 pairs each. Heat, cold, and drought are the best-sampled stresses; high light (6 species) has the most limited coverage (Fig. 1A). Our in-house stress experiments on *Brachypodium distachyon* (Fig. S1), *Selaginella moellendorffii* (Fig. S2) and *Klebsormidium nitens* (Fig. S3) produced visible, dose-dependent phenotypes confirming effective stress application across divergent plant lineages (Fig. 1B, C). As a first pass validation of the dataset, we obtained a set of known canonical marker genes for each stress type, and queried for their expression values across the dataset (Table S2). We observed that the canonical stress markers show consistent upregulation across species for heat (HSPs, HSFA2), cold (CBFs, CORs), drought (RD29A/B, RAB18), pathogen (PR1-3, WRKY33), and flooding (ADH1, PDC1, HRE1), validating that the cross-species DEG pipeline recapitulates known biology (Fig. 1D, orthogroups found in Supplementary Dataset 1). Interestingly, salt markers (SOS1, NHX1, HKT1) show wider spread centered near zero, consistent with their role as constitutive ion transporters rather than transcriptionally induced genes. Herbivory and heavy metal markers show positive but modest induction across fewer species, reflecting sparser experimental coverage and more lineage-specific responses (Fig 1D).

**Figure 1.**
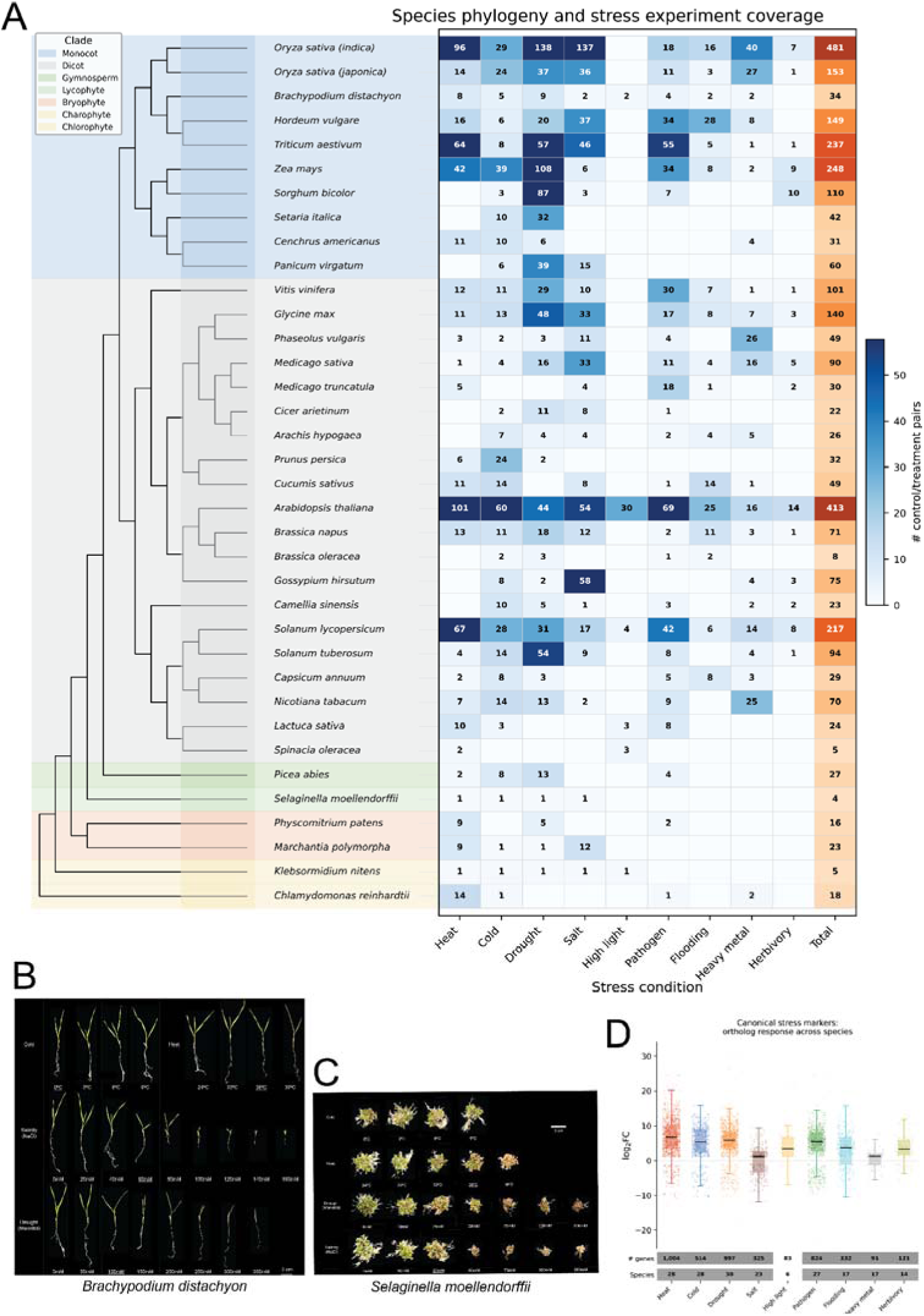
Kingdom-wide stress transcriptome atlas and marker gene validation. (A) Phylogenetic relationships of 36 species colored by clade (monocot, dicot, gymnosperm, lycophyte, bryophyte, charophyte, chlorophyte), coupled with a heatmap showing the number of control/treatment pairs per species per stress condition. Total column (orange) shows cumulative coverage per species. (B) Representative phenotypes of *Brachypodium distachyon* under stress treatments (heat, salt, drought), showing visible morphological responses at increasing treatment intensities. The underlined conditions were selected for RNA sequencing. (C) Representative phenotypes of Selaginella moellendorffii under stress treatments, demonstrating stress-induced morphological changes in a lycophyte. (D) Distribution of log2FC values for orthologs of GO-verified canonical stress marker genes across species, shown as stripplots with boxplot overlays per stress type. Marker genes were selected by intersecting curated literature candidates with TAIR experimental GO evidence (IMP, IDA, IEP, IGI codes), then mapped to orthologs via OrthoFinder. Annotation rows show the number of ortholog-experiment data points and the number of species with data per stress.

### Ancient gene families respond broadly but moderately; young families respond intensely but narrowly

We structured the dataset into hierarchical categories which allows for species, stress, organ, bioproject and direction (up- or down-regulated) querying (Fig. 2A), and subjected it to differential expression analysis with DeSeq2 (Love et al., 2014), which yielded ∼13,600,000 differentially expressed genes (DEGs, Supplementary Dataset 2). We observed that leaf, root, seedling and shoot were the most dominant organ categories across the various stresses (Fig. 2B). In order to understand how the conservation of stress DEGs were distributed across the phylogenetic lineage, we used OrthoFinder (Emms & Kelly, 2019) to group the genes from the 36 species into orthogroups (OGs). We next identified OGs in which at least one stress DEG from any species was present, and plotted the distribution of these stress OGs with respect to the number of species-specific stress DEGs observed, from which we calculated a conservation score (Fig. 2C, Table S3). We observed that the conservation score was bimodally distributed, pointing to two classes of OGs. The first class represents the vast majority (∼87%) of stress-responsive OGs that are lineage-specific, found in fewer than 10% of species. The second class represents ∼8,700 OGs (∼8.5%) that are universally conserved across >50% of species (Fig. 2C). This led us to question if different stresses would exhibit stress-specific DEG conservation. For each stress, we plotted the proportion of universal (CS > 50%), moderate (10-50%), and lineage-specific (< 10%) OGs (Fig. 2D). We observed that the proportion of universal OGs differed in each stress, with heat, cold, drought, pathogen and heavy metal showing significant proportions of universal OGs, suggesting that responses to these stresses were more highly conserved than flooding, herbivory and high light (Fig. 2D).

**Figure 2.**
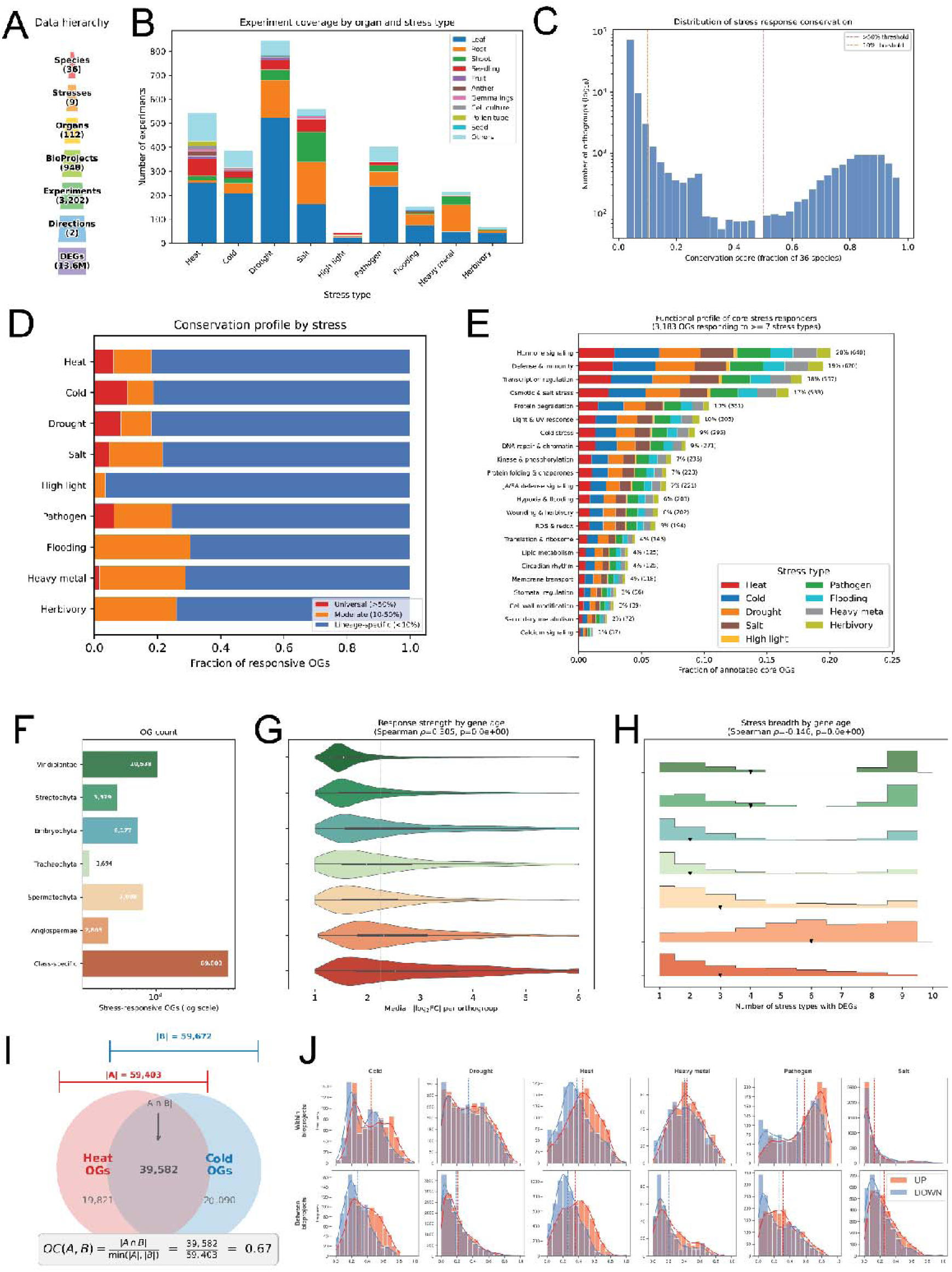
Gene family analysis of stress responses across the plant kingdom. (A) Data hierarchy of the kingdom-wide stress transcriptome atlas, spanning 36 species, 9 stress types, and 13.6 million DEG calls. (B) Experiment coverage by organ and stress type, showing number of unique species-BioProject-experiment combinations. (C) Distribution of orthogroup conservation scores, defined as the fraction of 36 species with at least one DEG in each orthogroup (log10-scaled y-axis). Dashed lines indicate thresholds for universal (>50%) and lineage-specific (<10%) classification. (D) Fraction of responsive orthogroups classified as universal, moderate, or lineage-specific for each stress type. (E) Functional profile of 3,183 core stress responders (orthogroups with DEGs in >=7 of 10 stresses), annotated using curated GO biological process categories mapped through Arabidopsis orthologs. Bar segments show the relative contribution of each stress type, weighted by species count. (F) Number of stress-responsive orthogroups per phylostratum, assigned by the most basal taxonomic group containing members of each orthogroup. (G) Response strength by gene family age, showing the distribution of median |log2FC| per orthogroup (clipped at 6) as violin plots per phylostratum.(H) Stress breadth by gene family age, showing the number of stress types each orthogroup responds to, displayed as ridgeline histograms per phylostratum. Triangles indicate medians. (I) Schematic of the overlap coefficient used to quantify shared stress-responsive orthogroups between stress pairs. (J) Distribution of overlap coefficients for up-regulated (red) and down-regulated (blue) DEGs within BioProjects (top row) and between BioProjects (bottom row) for six well-sampled stresses. Dashed lines indicate meanvalues.

Next, we wanted to identify core biological processes which were conserved across the various stresses. We first obtained orthogroups with DEGs in >=7 of 9 stresses, and annotated the orthogroup members using curated GO biological process categories mapped through *Arabidopsis* orthologs. We then calculated the summed proportion of each species and stress that were represented by a particular GoSlim biological process category. The barchart shows that core stress responders (active across >=7 stresses) are enriched for hormone signaling (20%), defense/immunity (19%), transcription regulation (18%), and osmotic/salt stress response (17%), representing the conserved stress signaling backbone of green plants (Fig. 2E).

To characterize the phylogenetic differences in stress response, we performed the phylostratigraphic analysis by categorising the OGs into phylogenetic clades based on the oldest lineage observed in that particular OG (Fig. 2F) (Table S4) (Domazet-Lošo et al., 2007). For each OG, we calculated the number of unique stresses represented by their stress-specific DEGs present in the OG, and then plotted the distribution of these OGs for each clade (Fig. 2H). We observed that there were two distinct periods of stress response evolution. A large number of highly conserved stress OGs (>= 7 stresses) were formed in the clade Viridiplantae, slowly decreasing as plants evolved towards Streptophyta, Embryophyta and Tracheophyta. During the evolution of the Angiosperms, a sudden large increase in the number of highly conserved stress OGs was again observed. This may be due to the superior evolutionary fitness of Angiosperms which enabled them to rapidly reproduce and adapt to various ecological habitats (Condamine et al., 2020). Furthermore, we observed that the ancient gene families (Viridiplantae/Streptophyta age) respond to more stress types (median 4) but with moderate fold changes (median |log_2_FC| = 1.5-1.7), while younger, class-specific families respond to fewer stresses (median 3) but with stronger induction (median |log_2_FC| = 2.5) (Fig. 2G and H). This suggests a division of labor: ancient gene families provide broadly deployed, moderate stress housekeeping, while younger families deliver intense, specialized responses.

The subsequent analyses in this study compare DEG lists that often differ substantially in size. To account for this, we adopted the overlap coefficient (OC), defined as the intersection size divided by the size of the smaller set (Fig. 2I). As an initial assessment of dataset consistency, we computed OC values within and between BioProjects for each stress and regulation direction (Fig. 2J). Within-BioProject OC values are consistently higher than between-BioProject values for all stresses except salt, confirming that experimental context strongly influences which genes are detected as differentially expressed. Upregulated DEGs also overlap more than downregulated ones in both settings, suggesting that stress activation programs are more conserved across experimental conditions than suppression programs.

### Stress response conservation is shaped by organ context, phylogenetic distance, and stress type

Most stress transcriptomics studies focus on a single organ, yet it is unclear whether stress responses are organ-specific or reflect a shared core program. To test this, we compared OC distributions measured within the same organ (organ-restricted) versus across different organs (cross-organ) for each stress and regulation direction (Fig. 3A). OC distribution within organs (Organ-restricted - pink) was always consistently higher than that measured across organs (Cross-organ - blue) (Fig. 3A), with the exception of salt stress. When compared against a shuffled distribution where OCs were computed additionally across stresses, the organ-restricted distributions consistently showed significant differences for most stresses, while the cross-organ distributiosns were significant mainly in the up-regulated DEGs (Fig. 3A, Top). Organ-restricted OC values were consistently higher than cross-organ values across most stresses, indicating that a substantial fraction of the stress response is organ-dependent. However, cross-organ OC values still exceeded the shuffled null distribution for most stresses, particularly for upregulated DEGs, demonstrating that a common transcriptional core is shared across tissues. Salt and drought were exceptions, with neither organ-restricted nor cross-organ means exceeding the shuffled distribution.

**Figure 3.**
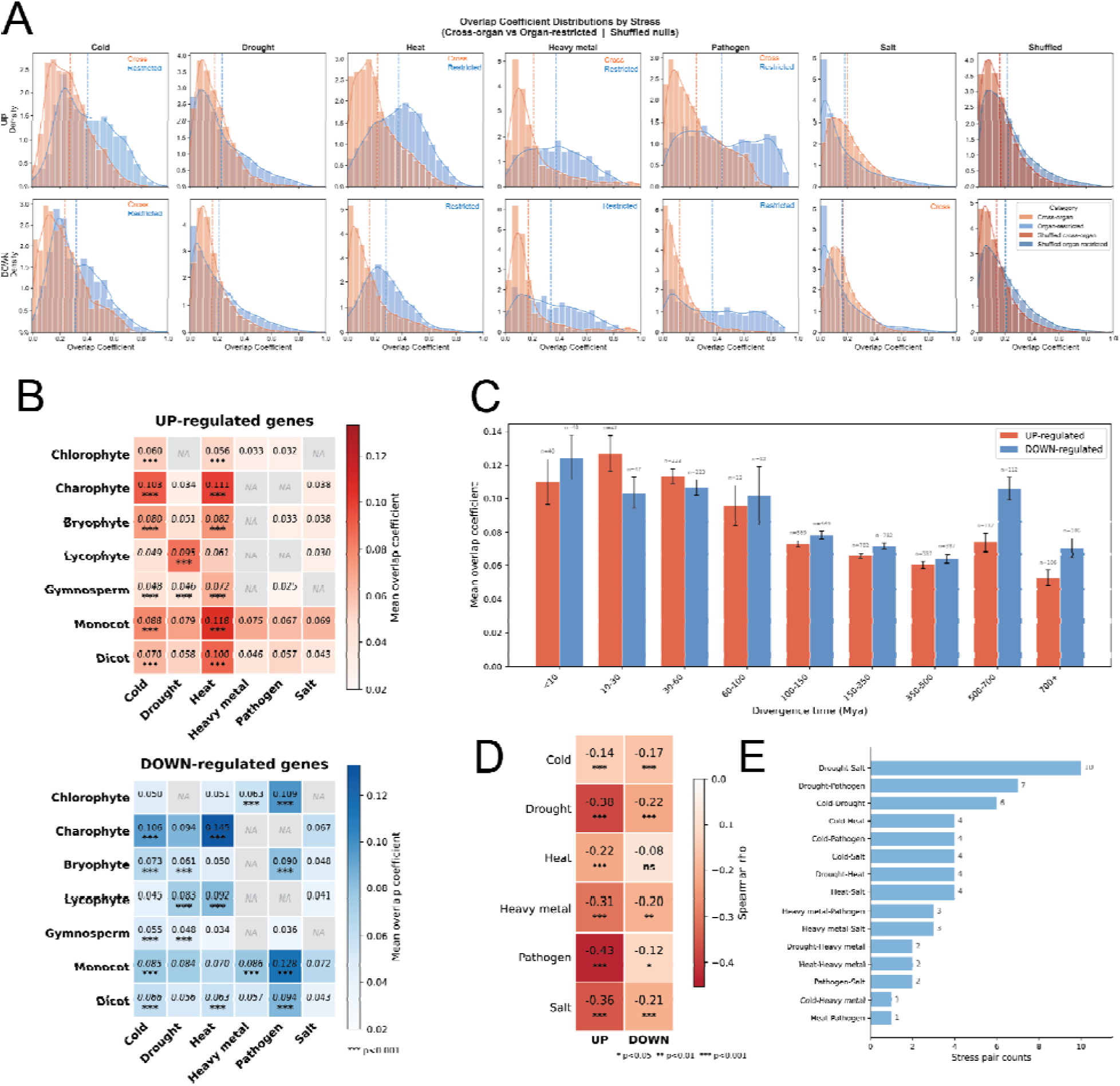
Overlap coefficients reveal organ-specific, clade-specific, and cross-stress similarities between stress responses across Viridiplantae. (A) Density distributions of overlap coefficients (OC) comparing stress-responsive orthogroup sets between conditions within the same organ (organ-restricted, blue) and across different organs (cross-organ, orange), shown per stress type (columns) and DEG direction (rows: UP, DOWN). Dashed vertical lines indicate meanOC values. Significance was assessed by Monte Carlo permutation (1,000 iterations). The rightmost column (“Shuffled”) pools all shuffled comparisons regardless of stress. (B) Heatmaps of mean cross-clade overlap coefficients for UP-regulated (top, red) and DOWN-regulated (bottom, blue) orthogroups across six stresses (columns) and seven plant clades (rows). Significance was assessed by Monte Carlo permutation (1,000 iterations). Cells marked NA indicate insufficient data. *** p < 0.001. (C) Mean overlap coefficient as a function of pairwise phylogenetic distance, binned into nine divergence time intervals, for UP-regulated (red) and DOWN-regulated (blue) orthogroups. Values were pooled across six stresses. Error bars indicate standard error of the mean; n values denote the number of species-pair x stress combinations per bin. (D) Heatmap of Spearman rank correlation coefficients (rho) between pairwise divergence time and mean OC, per stress type and DEG direction. * p < 0.05, ** p < 0.01, *** p < 0.001. (E) Number of species in which each stress pair showed significantly higher OC than expected by chance (permutation test, adjusted p < 0.05).

We next examined whether stress responses are conserved across phylogenetic clades. For each species, all DEGs were translated into one-to-one orthologs of every other species using OrthoFinder output, ensuring that no single species dictionary biased the comparison. OC values were then computed exclusively between species from different clades, pooled across species-specific dictionaries, and tested for significance against a cross-stress null distribution (Fig. 3B). Heat, cold and drought responses are significantly conserved across clades in both upregulated and downregulated DEGs, while heavy metal and pathogen responses are conserved only in downregulated DEGs. Salt responses do not reach significance in most cross-clade comparisons, suggesting a lineage-specific adaptation program.

To quantify how stress response conservation decays over evolutionary time, we converted clade-pair comparisons to divergence times using TimeTree (Kumar et al., 2022) and computed mean OC values at each time interval (Fig. 3C, Fig. S4). Downregulated genes show slightly higher OC than upregulated genes at most divergence bins, and upregulated genes decay faster than downregulated genes across all stresses. Spearman rank correlation confirmed significant negative relationships between divergence time and mean OC for all stress-direction combinations (Fig. 3D). Among individual stresses, pathogen shows the steepest decay for upregulated genes (rho = -0.43), consistent with the rapid evolution of plant immunity, while heat shows the weakest decay, consistent with the deep conservation of the heat shock response (Lindquist, 1986).

Finally, we asked whether certain stress pairs share transcriptional programs more than others. For each species, we computed OC values between DEG lists from different stresses within the same organ, tested significance by Monte Carlo permutation, and tallied the number of species showing significant overlap for each stress pair (Fig. 3E). Drought and salt show the highest overlap (significant in ∼30% of species), consistent with shared osmotic and ABA signalling. Drought also overlaps frequently with pathogen (7 species), suggesting crosstalk between dehydration and defence pathways. Heat and pathogen show the least overlap (1 species), indicating largely independent transcriptional programs.

### Conserved and stress-specific functional co-occurrence networks across the plant kingdom

To identify biological processes that are coordinately regulated under stress, we used Mercator (Lohse et al., 2014) to assign functional annotations to all genes across the 36 species. For each combination of species, stress and organ, we computed pathway enrichment scores summarising the relative enrichment of each functional bin among up- and down-regulated DEGs (Table S5). A direction bias, defined as the difference between the up and down enrichment scores, was then calculated for each bin across groups of interest. This analysis reveals a clear split between co-upregulated defence processes and co-downregulated growth processes (Fig. 4A). Protein quality control, glutathione-based redox regulation, galactose metabolism and amino acid degradation are consistently upregulated across all six stresses, while cell wall proteins, pectin, microtubular network, photophosphorylation and brassinosteroid signalling are consistently downregulated. Protein quality control is the most up-biased bin across all stresses; cell wall proteins is the most down-biased. This growth-defence tradeoff is conserved from chlorophytes to angiosperms to a varying degree (Fig. S5).

**Figure 4.**
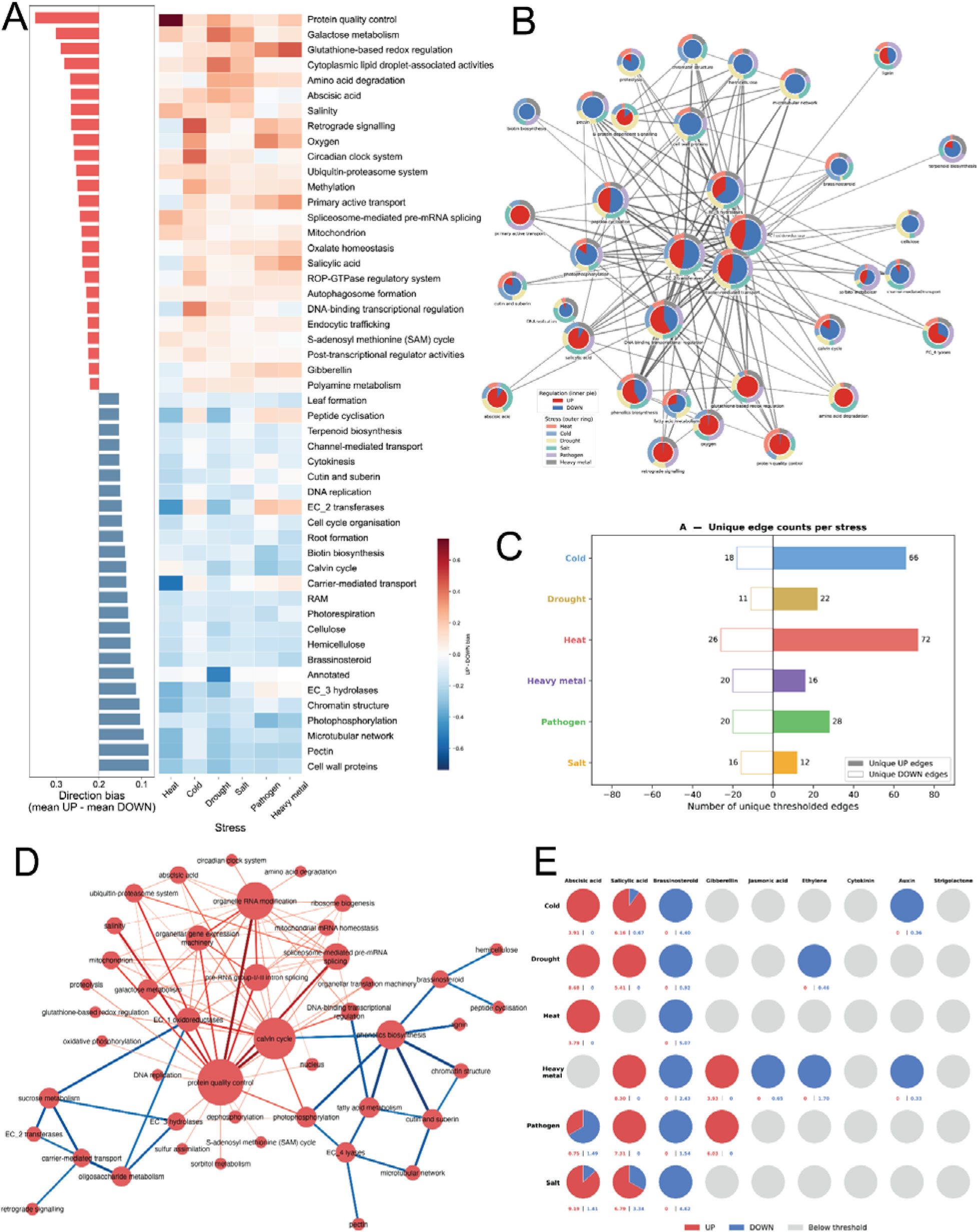
Functional pathway co-occurrence reveals a conserved growth-defense tradeoff and stress-specific transcriptional rewiring across 36 plant species. A) Direction bias of MapMan Level 1 bins across heat, cold, drought, salt, pathogen, heavy metal. Left: mean direction bias (UP - DOWN enrichment score) averaged across all stresses and 36 species. Right: per-stress direction bias heatmap. Bins are ranked from most UP-biased (protein quality control, top) to most DOWN-biased (cell wall proteins, bottom). Red indicates co-upregulation, blue indicates co-downregulation. The direction bias represents the difference between the fraction of experiments where a bin’s DEGs are upregulated versus downregulated. B) Mercator Level 1 functional co-regulation network across abiotic and biotic stress conditions. Each node represents a Mercator Level 1 functional bin present in more than three stress types. Node size is proportional to total weighted degree. Node colouring represents the proportion of total weighted degree attributable to up- (red) versus down- (blue) regulation, displayed as a pie chart. Edges connect bin pairs that co-occur in more than three stress-type networks. Ring colours indicate the contributing stress type. C) Number of functional co-regulation network stress-unique edges. Solid bars: unique UP edges (co-upregulated only in that stress). Hatched bars: unique DOWN edges (co-downregulated only in that stress). D) Heat-specific functional co-regulation network. Nodes represent Mercator Level 1 functional bin categories; node size is proportional to total weighted degree within that stress’s unique network. Edge colour indicates direction of co-regulation: red = co-upregulated, blue = co-downregulated; edge width and colour intensity are proportional to edge weight. E) Pie charts show the balance of UP- (red) and DOWN-regulated (blue) functional co-regulation edge weights for the first neighbours of each phytohormone. Numbers below each pie indicate the summed UP (left) and DOWN (right) edge weights above threshold. Grey circles indicate hormones present in the network but with no edges above threshold in that organ.

Stress responses are not isolated pathway activations but involve coordinated shifts across multiple cellular processes (Lee et al., 2025). To understand how these processes are wired together, we constructed functional co-occurrence networks from the pathway enrichment results. For each DEG list, we built a binary bin-by-bin matrix recording which functional categories co-occurred among up- or downregulated genes, then averaged these matrices across all experiments within a group (Table S6). The resulting network for the six stresses, filtered to bins present in at least four stresses, reveals a conserved core of co-regulated processes alongside stress-specific connections (Fig. 4B). Several co-regulatory relationships are maintained regardless of stress type: cell wall proteins, pectin and microtubular network form a tightly connected downregulated cluster. Hormone bins show a conserved split, with ABA and salicylic acid embedded in the upregulated subnetwork and brassinosteroid in the downregulated one. The stress-type composition of each edge, shown as ring colours around each node, reveals that most high-degree nodes receive contributions from all six stresses, confirming that the core wiring of the stress response is shared rather than stress-specific.

To identify stress-specific rewiring, we counted the number of unique edges per stress after removing edges shared across all six stresses (Fig. 4C). Cold and heat stand out with the most unique upregulated edges (66 and 72 respectively), indicating that both stresses activate distinctive sets of co-regulated processes. However, they differ in their downregulated signatures: heat has 26 unique downregulated edges compared to only 18 for cold, consistent with the broader suppression of cellular activity observed under heat throughout our analyses (Fig. 4A). Pathogen and drought show intermediate numbers of unique edges in both directions, while salt and heavy metal have the fewest, suggesting their transcriptional responses overlap more extensively with other stresses, or that the breadth of transcriptional responses to these stresses are smaller (Fig. 4C). The biological function of the stress-specific co-regulated functions align well with our knowledge of the stress. Heat uniquely links protein quality control to organelle RNA modification and pre-mRNA splicing (Fig. 4D), which are known to be affected in heat (Ling et al., 2021). Cold stress is organised around the circadian clock system as a central hub (Bieniawska et al., 2008; M. A. Dong et al., 2011), connecting to retrograde signalling and transcriptional regulation (Fig. S6 for other stresses). Drought centres on cytoplasmic lipid droplet-associated activities (Vries & Ischebeck, 2020), linking to ABA (Muhammad Aslam et al., 2022), galactose metabolism and amino acid degradation. Salt connects cutin and suberin to fatty acid metabolism, consistent with Casparian strip reinforcement (Chen et al., 2011). Pathogen response forms a defence cluster around phenolics biosynthesis and lignin (Miedes et al., 2014). Heavy metal response is organised around sulfur assimilation, connecting to glutathione-based redox regulation and carrier-mediated transport (D. Mendoza-Cózatl et al., 2005).

Finally, to better understand the role of hormones in stress responses, we extracted the subnetworks of phytohormone annotated bins (Fig. 4E). Brassinosteroid is consistently downregulated across all stresses, supporting its role in orchestrating the growth side of the growth-defence tradeoff (Nolan et al., 2020). ABA and salicylic acid are predominantly upregulated, consistent with their established roles in stress signalling (Yang et al., 2023). Different stresses activate different combinations of hormone subnetworks: ABA dominates drought and salt, salicylic acid is broadly active across both abiotic and biotic stresses, and gibberellin is upregulated specifically under pathogen and heavy metal. Cytokinin remains below the responsiveness threshold in all stresses. To validate these hormone-stress associations against existing knowledge, we compared our transcriptomic direction bias scores with the number of literature entries linking each hormone to each stress in the PlantConnectome knowledge graph (4.8 million edges, predominantly Arabidopsis-derived; Fig. S7) (S. C. Lim et al., 2025). The knowledge graph is heavily skewed toward pathogen-hormone interactions (JA-Pathogen: 1,147 entries; Ethylene-Pathogen: 761), while abiotic stress-hormone links are sparse for most hormones. Our cross-kingdom data reveal two notably underappreciated patterns. First, salicylic acid, with only 1-3 literature entries for abiotic stresses, emerges as broadly upregulated under cold (+0.15), pathogen (+0.21) and heavy metal (+0.29), making it one of the most broadly responsive stress hormones across the kingdom. Second, brassinosteroid is consistently and strongly downregulated across all six stresses (up to -0.22 under heat), yet has only 10-40 literature entries per abiotic stress. Given that BR suppression tracks closely with the growth-defence tradeoff observed throughout our analyses (Fig. 4A), its transcriptional role as a conserved mediator of growth suppression under stress appears substantially underexplored. These discrepancies highlight the value of unbiased, cross-kingdom expression analysis for identifying gaps in current knowledge.

### Conserved stress co-expression modules undergo regulatory subfunctionalization through duplication across the plant kingdom

To characterize the modular organization of stress transcriptional programs, we constructed gene co-expression networks using the collected RNA-seq data using TEA-GCN (P. K. Lim et al., 2026) and applied Louvain community detection to identify stress-responsive modules. Modules were compared across species using the overlap coefficient on their orthogroup content, and conserved pairs (OC >= 0.5) were assembled into a cross-species module network. 31 out of 36 species produced networks with sufficient z-score range for downstream analysis (Fig. S8, Supplementary Dataset 3). For each species and stress type, we extracted the subnetwork connecting differentially expressed genes (z >= 1.7) and applied Louvain community detection to identify stress-responsive co-expression modules (Blondel et al., 2008). To assess conservation, modules were compared across species using the overlap coefficient on their orthogroup content, and conserved module pairs (OC >= 0.5) were assembled into a cross-species module network and clustered using a second round of Louvain community detection to identify groups of functionally related modules (Supplementary Dataset 4).

To illustrate the outcome of this analysis, we focus on heat stress (Fig. 5A). Heat stress modules are conserved across more than 20 species spanning multiple lineages (similar modules connected by gray lines), though predominantly in flowering plants. We identified five groups of conserved modules (Fig. 5A, indicated by different colors), indicating that heat responses use five distinctive transcriptional programs. Notably, Arabidopsis modules 4 and 5 are connected by paralog edges (red lines), showing that these two modules share substantial orthogroup content, indicating that they represent duplicated modules within the same genome, with shared genes regulated in divergent directions under heat stress.

**Figure 5.**
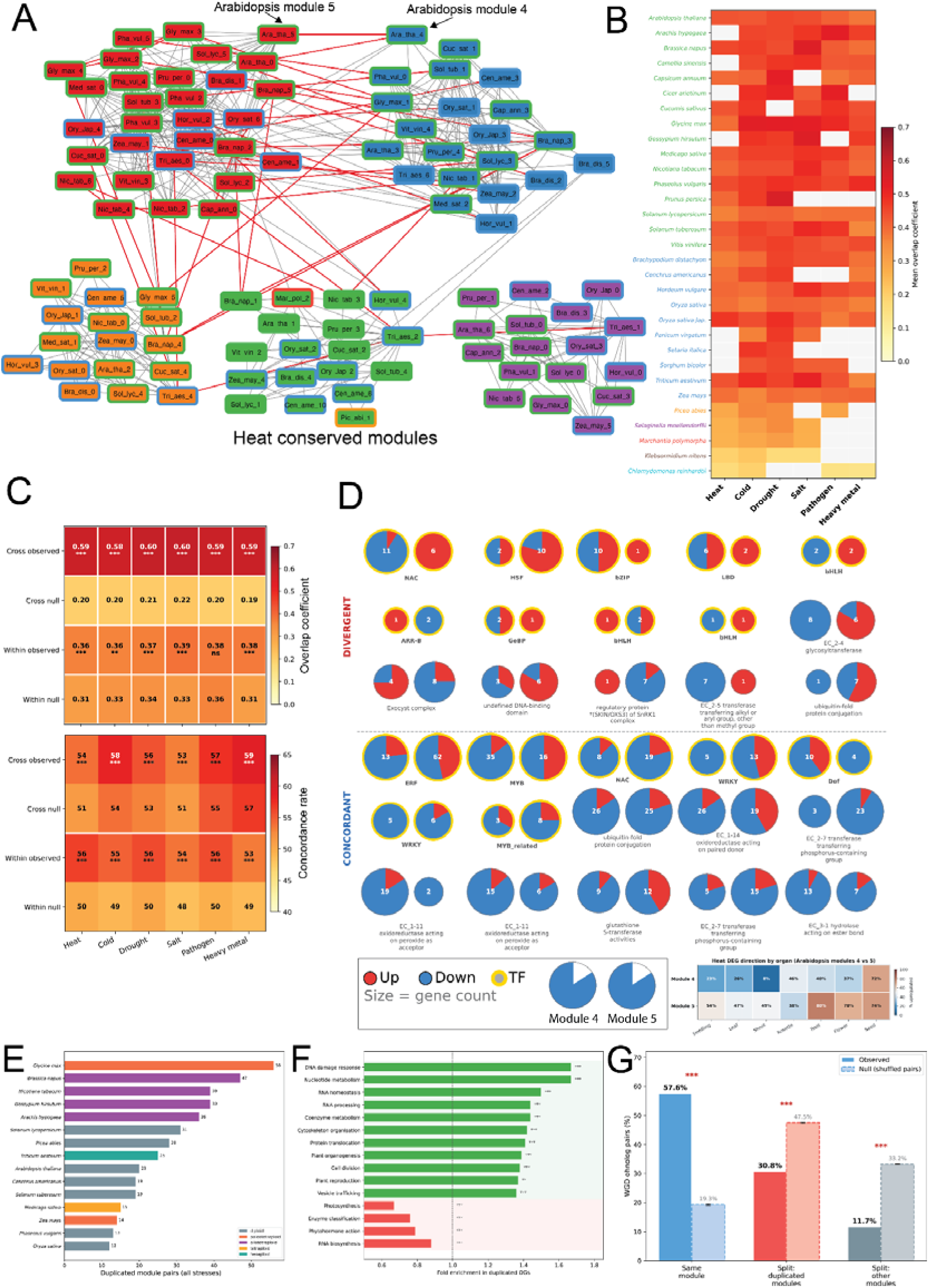
Conserved stress co-expression modules across the plant kingdom reveal regulatory subfunctionalization through module duplication. (A) Cytoscape network of heat stress co-expression modules conserved across species (overlap coefficient >= 0.5, >= 10 shared orthogroups). Nodes represent species-specific modules; edges represent orthogroup overlap between modules. Node fill color indicates network cluster assignment; node border color indicates lineage. Red edges connect duplicated module pairs within the same species (OC >= 0.3). Arabidopsis modules 4 and 5 and their cross-species orthologs are highlighted. (B) Cross-species conservation heatmap showing mean best overlap coefficient per module to all other species, for 31 species across 6 stress types. Species are ordered by lineage (color-coded: green = dicot, blue = monocot, orange = gymnosperm, purple = lycophyte, red = bryophyte, brown = charophyte, cyan = chlorophyte). White cells indicate missing data (stress not profiled in that species). (C) Overlap coefficient (top) and direction concordance rate (bottom) for cross-species conserved module pairs (OC >= 0.5, n = 4,503) and within-species duplicated pairs (OC >= 0.3, n = 498), compared against a null model with shuffled gene-to-orthogroup assignments (100 permutations). White line separates cross-species from within-species rows. Wilcoxon signed-rank test (* p < 0.05, ** p < 0.01, *** p < 0.001). (D) Shared orthogroups between Arabidopsis heat modules 4 (81% downregulated) and 5 (46% upregulated). Each pair of pies represents one orthogroup: left pie = module 4, right pie = module 5. Pie size reflects gene copy number; red/blue slices show up/down proportions across all copies. Yellow borders indicate transcription factor families. Top section: divergent OGs (opposite majority direction); bottom section: concordant OGs (same direction). 36 representative OGs shown of 585 total shared. The heatmap below shows the percentage of upregulated genes across organs in the two modules. (E) Within-species module duplication across top 15 species by pair count shown. Bar color indicates ploidy level. (F) Functional enrichment of orthogroups found in duplicated module pairs versus all annotated orthogroups (Fisher’s exact test, BH correction). (G) Location of WGD ohnolog pairs relative to stress modules (10 species with PLAZA dicots/monocots 5.0 syntenic anchor data). For ohnolog pairs where both genes are in modules (n = 378,021 pair-stress instances): “Same module” = both copies in the same module; “Split: duplicated modules” = one copy in each module of a duplicated pair; “Split: other modules” = copies in unrelated modules. Dashed bars show null expectation from 100 permutations of ohnolog pair assignments. Significance by z-test against permutation distribution (*** p < 0.001).

The conservation pattern is consistent across all six stress types (mean OC 0.3-0.5), with strongest conservation among angiosperms and decreasing signal with phylogenetic distance (Fig. 5B). The uniformity across stresses suggests that modular transcriptional architecture is ancient and largely stress-independent. Null model comparison (100 permutations of gene-to-orthogroup assignments) confirmed that cross-species module conservation is 3-4x above chance for all stresses while within-species duplicated module pairs showed Jaccard indices only marginally above null, consistent with modules from the same genome drawing from a shared orthogroup pool (Fig. 5C, Fig. S9). Direction concordance is modestly but significantly elevated above null (+2-6 percentage points), indicating that most concordance reflects baseline expression proportions rather than orthogroup-specific regulatory conservation.

Closer inspection of the 585 shared orthogroups between Arabidopsis modules 4 and 5 reveals the logic of this duplication (Fig. 5D). Stress-responsive transcription factor families (NAC, WRKY, bZIP, HSF) show divergent regulation between the two modules, being downregulated in module 4 and upregulated in module 5, consistent with module 4 representing a growth program suppressed under heat and module 5 an active defence response. Enzyme families such as oxidoreductases and transferases maintain concordant regulation in both modules, indicating that regulatory divergence is driven by transcription factors while shared downstream targets respond in the same direction. Many of the shared orthogroups are represented by multiple paralogs within each module, and the up/down ratios across these copies are not uniform, indicating that regulatory fine-tuning also operates at the level of individual gene copies within orthogroups.

Within-species module duplication was detected in 28 of 31 species with sufficient data, totaling 504 duplicated module pairs across all stresses (Fig. 5E). Polyploid species harbored the most duplicated pairs, led by the paleotetraploid *Glycine max* (56 pairs) and the allotetraploid *Brassica napus* (47 pairs), followed by the allotetraploids *Nicotiana tabacum* and *Gossypium hirsutum* (39 pairs each). However, diploid species also showed substantial module duplication, including *Picea abies* (28 pairs) and *Solanum lycopersicum* (31 pairs), indicating that within-species module duplication is not solely a consequence of whole-genome duplication but a general feature of stress-responsive transcriptional networks.

Functionally, orthogroups shared between duplicated module pairs are enriched for cellular infrastructure, including DNA damage response (1.67x), nucleotide metabolism (1.67x), RNA homeostasis (1.50x), RNA processing (1.44x), cytoskeleton organisation (1.42x), cell division (1.38x) and vesicle trafficking (1.36x), while depleted for photosynthesis (0.67x), phytohormone action (0.79x) and transcription factors (0.88x; Fig. 5F). This depletion of transcription factor orthogroups among shared genes suggests that duplicated modules diverge through different regulatory repertoires while retaining shared downstream machinery.

Tracking WGD ohnolog pairs across 10 species (PLAZA dicots/monocots 5.0) (Van Bel et al., 2022) revealed that ohnologs overwhelmingly co-localize in the same module (57.6% vs 19.3% null) and avoid splitting into unrelated modules (11.7% vs 33.2% null; Fig. 5G). The fraction landing in duplicated module pairs (30.8%) was below null expectation (47.5%), indicating that WGD is not the primary driver of module duplication but rather reinforces existing co-regulatory relationships.

### Cis-regulatory sequence features predict stress-responsive genes across the plant kingdom

Our stress atlas provides an unprecedented opportunity to ask a fundamental question in plant biology: is stress-responsive gene expression encoded in the DNA and RNA sequence/structure itself, and if so, what are the sequence features that determine it? We explored several models that can predict gene expression from DNA or RNA sequence (Fig. 6A): a deep convolutional neural network (CNN) baseline (based on (Peleke et al., 2024), and two foundation models: PlantCAD2 (Zhai et al., 2025), a Mamba/Caduceus DNA language model pre-trained on 65 plant genomes (88M parameters), and PlantRNA-FM, an ESM-based RNA language model pre-trained on transcriptomes from 1,124 plant species (33M parameters) (Yu et al., 2024). All models were trained on DNA and RNA sequences from 25 plant species spanning angiosperms, gymnosperms, bryophytes and green algae. For each stress and regulation direction, genes were labeled as stress-responsive or non-responsive based on their DEG frequency quartile across experiments in the Kingdom Stress Atlas. Model weights and instructions on how to run them are available in Supplementary Dataset 5.

**Figure 6.**
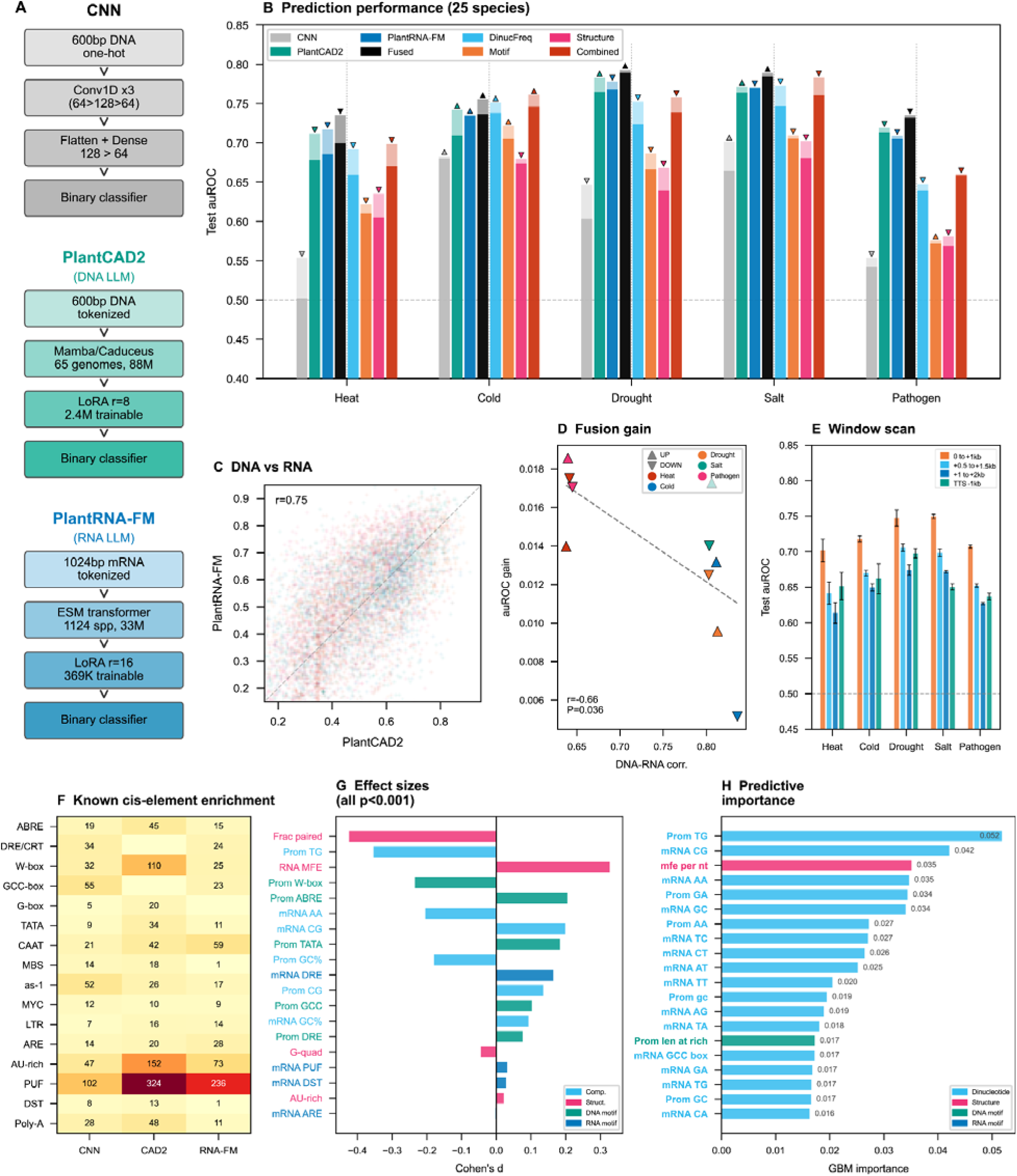
Predicting stress-responsive genes from DNA and RNA sequence across 25 plant species. (A) Architectures of the three sequence-based models. (B) Test auROC across five abiotic and biotic stresses for eight models, comprising four sequence-based models (CNN, PlantCAD2, PlantRNA-FM, and their late fusion), four interpretable feature-based models using gradient-boosted machines (GBM) trained on dinucleotide frequencies (DinucFreq, 34 features), known and discovered cis-element counts (Motif, ∼92 features), RNA secondary structure descriptors (Structure, 8 features), and all features combined (Combined, ∼134 features). For each model, the faded bar shows the higher-performing direction (up- or down-regulated, indicated by triangle marker) and the solid bar shows the lower-performing direction. (C) Per-gene prediction comparison between PlantCAD2 and PlantRNA-FM across all test genes, colored by stress (r = 0.75). Each point represents one gene scored by both models. (D) Fusion gain (improvement in auROC from averaging DNA and RNA predictions over the better single model) as a function of DNA-RNA prediction correlation per stress. Triangles indicate regulation direction (up/down). (E) PlantRNA-FM performance when retrained on different 1 kb transcript windows: 5’ end (0 to +1 kb from TSS), mid-transcript (+0.5 to +1.5 kb), deep transcript (+1 to +2 kb), and 3’ end (last 1 kb before TTS). Error bars indicate the range between up- and down-regulated models. (F) Known cis-regulatory element enrichment in the top 200 attribution-derived motif sequences per stress and direction, summed across all five stresses. Columns show hits from CNN (integrated gradients), PlantCAD2 (*in silico* mutagenesis), and PlantRNA-FM (gradient x input) attributions. (G) Cohen’s d effect sizes for sequence and structure features between stress-responsive and non-responsive genes, averaged across all stresses and both regulation directions. All displayed features are significant after Bonferroni correction (P < 0.001). Feature categories: sequence composition (cyan), RNA secondary structure (magenta), DNA cis-elements (teal), RNA regulatory elements (blue). (H) Mean GBM feature importance (Gini importance) from the combined interpretable model, averaged across all stresses and directions. Feature categories colored as in (G). Values indicate mean importance scores.

The CNN baseline achieved a mean test auROC of 0.613, which, while above chance, was substantially lower than either foundation model across all five stresses (Fig. 6B). PlantCAD2 and PlantRNA-FM performed comparably (mean auROC 0.736 and 0.737, respectively), despite operating on DNA and mRNA sequence, suggesting that both molecule types carry similar amounts of regulatory information relevant to stress responsiveness. Per-gene predictions from the DNA and RNA models were moderately correlated (r = 0.75; Fig. 6C), indicating that the two models captured overlapping but partially distinct regulatory signals. We fused the DNA and RNA models by simple averaging of predicted probabilities per gene, and observed a modest increase in performance of the fused model (Fig 6B, black bars). This improvement was inversely correlated with inter-model agreement (r = -0.66, P = 0.036; Fig. 6D): stresses where DNA and RNA predictions diverged most (Heat, Pathogen; correlation ∼0.64) benefited most from fusion (gain up to 0.019 auROC), while stresses with higher inter-model correlation (Drought, Salt; correlation ∼0.81-0.84) showed smaller but still consistent gains.

To better understand where in the mRNA molecule the regulatory signal resides, we retrained PlantRNA-FM independently on four 1 kb transcript windows (Fig. 6E). The 5’ region (0 to +1 kb from TSS), encompassing the 5’ UTR and CDS start, carried the strongest signal (mean auROC 0.725). Performance decayed toward the mid-transcript (+0.5 to +1.5 kb: 0.674; +1 to +2 kb: 0.647), but the 3’ end (last 1 kb before TTS) retained notable predictive power (0.660), suggesting that regulatory elements near the poly(A) site also contribute to stress-responsive gene identity. This pattern was consistent across all five stresses.

To identify potential cis-regulatory motifs that the models had learned, we performed attribution analysis using integrated gradients for the CNN, in silico mutagenesis for PlantCAD2, and gradient-times-input for PlantRNA-FM (Table S7). From the top 200 highest-attribution sequences per stress and direction, we extracted recurring k-mers (k=8) and scanned them for matches to 16 known plant cis-regulatory elements (Fig. 6F). PUF-binding sites were the most frequently discovered element across all three models (102 hits in CNN, 324 in PlantCAD2, 236 in PlantRNA-FM), consistent with the role of Pumilio/FBF proteins in mRNA stability regulation under stress (Huh, 2021). AU-rich elements, which target transcripts for rapid degradation (Gutiérrez et al., 2002), were enriched particularly in PlantCAD2 attributions (152 hits). W-box motifs (WRKY transcription factor binding sites) were strongly enriched in PlantCAD2 (110 hits) but rare in CNN and PlantRNA-FM, suggesting that the DNA language model is particularly sensitive to transcription factor binding patterns. Conversely, GCC-box motifs (ethylene-responsive element binding) were primarily discovered by the CNN (55 hits), indicating that different model architectures capture complementary aspects of the cis-regulatory code.

Given the large training dataset and the motifs discovered through attribution analysis, we explored whether these motifs, together with other sequence features: dinucleotide frequencies, GC content, and RNA secondary structure descriptors (minimum free energy, fraction of paired bases, G-quadruplex potential, AU-rich runs), could be used to build simpler, more easily interpretable models. We trained gradient-boosted machines (GBMs) on three feature groups: dinucleotide frequencies (32 features), motif counts from known and discovered elements (∼92 features), and RNA structure descriptors (8 features), as well as all features combined (∼134 features) (Supplementary Dataset 6). Surprisingly, the combined interpretable model achieved a mean auROC of 0.723 (Fig. 6B, rightmost bar), approaching the performance of the foundation models (0.736-0.737). Dinucleotide frequencies alone (0.712) outperformed motif counts (0.657) and RNA structure features (0.643), indicating that broad sequence composition is more predictive than any individual regulatory element. Down-regulated genes were generally easier to predict than up-regulated genes across all model types (Fig. 6B, most triangles point downwards).

Statistical comparison of sequence features revealed that the largest effects comprised RNA secondary structure: stress-responsive mRNAs had fewer paired bases (Cohen’s d = - 0.42) and higher minimum free energy per nucleotide (d = +0.33), consistent with less stable structures enabling faster translational response and turnover (Table S8). Promoter TG dinucleotide frequency was strongly depleted in responsive genes (d = -0.35), while mRNA CG dinucleotide content was elevated (d = +0.20). Among known cis-elements, ABRE (ABA-responsive element) showed the strongest enrichment in stress-responsive promoters (d = +0.20), consistent with the central role of abscisic acid signaling in abiotic stress responses (Yamaguchi-Shinozaki & Shinozaki, 2005). Notably, RNA structure effect sizes exceeded those of individual motifs, underscoring the importance of post-transcriptional regulation. GBM feature importance analysis confirmed that dinucleotide composition features dominated prediction (Fig. 6H, Table S9). Promoter TG dinucleotide was the single most predictive feature (Gini importance 0.052), followed by mRNA CG dinucleotide (0.042) and RNA minimum free energy (0.035). Known cis-elements (ABRE, DRE, GCC-box, W-box) appeared among the top 20 features but with individually lower importance, suggesting that the cis-regulatory code for stress responsiveness is distributed across many sequence features rather than concentrated in a few master regulatory elements.

## Discussion

We constructed a cross-kingdom stress transcriptome atlas spanning 36 Viridiplantae species and nine stress types, providing the broadest comparative view of plant stress gene expression to date. The atlas extends well beyond previous cross-species stress comparisons, which have typically been limited to two to five species within a single lineage (Y. Dong et al., 2023; Hartmann et al., 2022) or a single stress across a modest phylogenetic range. By combining reanalyzed public data with targeted in-house experiments on underrepresented lineages, and by manually curating all control-treatment pairs, we aimed to balance taxonomic breadth with analytical rigour.

One of the clearest patterns in the atlas is the relationship between gene family age and stress response profile. Ancient gene families, dating to the base of Viridiplantae or Streptophyta, respond to more stress types but with moderate fold changes, while younger, class-specific families respond to fewer stresses but with greater intensity (Fig. 2G-H). This pattern is consistent with a model in which ancient families form a broadly deployed stress housekeeping layer, while lineage-specific families provide intense, specialized responses tailored to the ecological challenges faced by particular clades. A similar logic has been described for immune gene evolution in animals, where conserved innate immunity coexists with rapidly evolving adaptive components (Buchmann, 2014), and our findings suggest an analogous architecture in plant stress biology.

Stress transcriptome similarity decays with phylogenetic distance, as expected, but the rate of decay differs between upregulated and downregulated genes. Upregulated genes show steeper decay across all stresses, while downregulated gene sets are more conserved even at deep divergence times (Fig. 3B-C). One interpretation is that the suppression of growth-related processes under stress (photosynthesis, cell wall biosynthesis, cell division) is constrained by the limited number of ways a plant can shut down growth, whereas the activation of defence programs is more evolutionarily labile because species face different biotic and abiotic challenges. This is consistent with our finding that pathogen-responsive upregulated genes show the steepest decay with divergence time, reflecting the well-documented rapid evolution of plant immunity (Jones & Dangl, 2006). Heat, by contrast, shows the weakest decay, consistent with the deep conservation of the heat shock response across all domains of life (Lindquist, 1986).

The co-occurrence network analysis reveals a strikingly consistent pattern across all six well-sampled stresses: protein quality control, redox regulation and amino acid degradation are co-upregulated, while cell wall biosynthesis, photosynthesis and brassinosteroid signalling are co-downregulated (Fig. 4A-B). This growth-defence tradeoff is maintained from chlorophytes to angiosperms, suggesting it reflects a fundamental constraint on resource allocation rather than a derived regulatory circuit. Growth-defence tradeoffs have been extensively studied in individual species, particularly in the context of jasmonate and salicylate signalling in Arabidopsis (Huot et al., 2014; Karasov et al., 2017; Züst & Agrawal, 2017). Our cross-kingdom data suggest that the tradeoff operates at a more general level than hormone-specific antagonism: it is visible in the coordinated suppression of cell wall, cytoskeleton and brassinosteroid pathways regardless of the stress type or the species. Also, the primacy of protein quality control as a key stress regulated hub across stresses (Fig. 4A) and clades (Fig. S5) suggests that it is a general and ancient response to stress. Indeed, hormone-independent stress responses mediated by the global decrease in translational efficiency have been shown previously (Koh et al., 2021, 2022, 2023). Alongside this universal pattern, each stress has a distinctive co-occurrence signature. The unique association of circadian clock components with cold stress, lipid droplets with drought, and sulfur assimilation with heavy metal stress are consistent with known biology (Espinoza et al., 2010; Gidda et al., 2016; D. G. Mendoza-Cózatl et al., 2011) but have not previously been shown to be conserved across dozens of species simultaneously. The accuracy by which the co-occurrence networks were able to recapitulate known stress physiology suggests that it shows significant potential in revealing regulatory patterns in other stresses and species. Comparison with the PlantConnectome knowledge graph revealed that the literature is heavily biased toward pathogen-hormone interactions, while the broad abiotic stress responsiveness of salicylic acid and the consistent downregulation of brassinosteroid across stresses are underrepresented. This illustrates the value of unbiased, cross-kingdom transcriptomic analysis for identifying gaps in current knowledge.

The conservation of stress co-expression modules across species extends beyond simple gene-level overlap. Modules sharing orthogroup content are present across 20 or more species for well-sampled stresses, and within-species module duplication is widespread, occurring in 28 of 31 species (Fig. 5). Duplicated module pairs are enriched for cellular infrastructure genes (RNA processing, vesicle trafficking, cell division) and depleted for stress effectors and transcription factors, suggesting that it is the regulatory scaffolding of the stress response, rather than its effector components, that undergoes subfunctionalization. The finding that whole-genome duplicate ohnologs are preferentially retained within the same module, rather than split across duplicated modules, has implications for models of post-WGD gene fate. Rather than driving regulatory divergence, WGD appears to reinforce existing co-regulatory relationships, at least for stress-responsive genes. This is consistent with our previous findings (Ruprecht et al., 2016), and the gene balance hypothesis (Birchler & Veitia, 2012), which predicts that dosage-sensitive genes are preferentially retained as duplicates. The observation that this pattern holds across 10 species with available syntenic data suggests it is a general feature of plant genome evolution.

The ability to predict stress-responsive genes from DNA and RNA sequence alone at notable accuracy has practical implications for non-model species (Fig. 6). The fact that dinucleotide composition outperforms individual cis-regulatory element counts as a predictive feature suggests that the sequence signature of stress responsiveness is distributed across many weak signals. This is consistent with recent findings from large-scale promoter studies in animals, where regulatory information is encoded in the overall sequence composition of promoters rather than in discrete motifs alone (Vaishnav et al., 2022). Furthermore, a similar pattern was observed for a cold stress-related study (Meng et al., 2021), suggesting that stress responses are governed by a joint effort of small sequence/structure features, rather than concentrated in a few master regulatory elements.

## Materials and Methods

### Stress transcriptome dataset assembly (Figure 1)

RNA-seq data were obtained from two sources: (1) publicly available datasets from SRA/ENA, reanalyzed using LSTRAP-Cloud (Tan et al., 2020) with Kallisto (Bray et al., 2016) for transcript quantification, and (2) in-house experiments on B*rachypodium distachyon, Selaginella moellendorffii,* and *Klebsormidium nitens*, quantified directly with Kallisto. Samples passing quality control (n_pseudoaligned >= 2 million reads and an adaptive pseudoalignment percentage cutoff) were retained. Control-treatment pairs were manually curated for each species, stress, and BioProject. Differential expression was computed using DESeq2 1.42.1 (Love et al., 2014), with DEGs defined as genes with |log_2_FC| > 1 and adjusted p-value < 0.05. The final dataset comprises 36 species spanning monocots, dicots, gymnosperms, lycophytes, bryophytes, charophytes, and chlorophytes, covering 9 stress types (heat, cold, drought, salt, high light, pathogen, flooding, heavy metal, and herbivory).

### Stress experiments for Brachypodium distachyon, Selaginella moellendorffii and Klebsormidium nitens (Figure 1)

Three species representing divergent plant lineages were subjected to abiotic stress treatments: *Brachypodium distachyon* (Bd21-3), *Selaginella moellendorffii*, and *Klebsormidium nitens* (NIES-2285). *B. distachyon* was grown on full-strength modified Hoagland solution agar at 24℃, 16/8 hours light/dark, 100 umol m-2 s-1. *S. moellendorffii* was cultivated on half-strength MS agar with vitamins at 24 ℃, 16/8 h L/D, 40 umol m-2 s-1. *K. nitens* was propagated in modified 3NBBM-V liquid medium under continuous shaking at 100 rpm, 40 umol m-2 s-1. All species were maintained in AR-95L Percival growth chambers at 24 ℃, 50% relative humidity.

For each species, a scoping experiment was performed across a severity gradient for heat, cold, salt and osmotic/drought stress to select optimal treatment intensities for RNA-seq. Ages for the respective species at the point of treatment were 10 days (*Brachypodium*), 8 days (*Selaginella*), OD 0.3 (*Klebsormidium*) respectively. Environmental stresses (heat, cold) were applied for 1 day, while media stresses (salt, osmotic/drought) were applied for the full growth period. Phenotypic responses were quantified using ImageJ (shoot length, root length, leaf count, surface area for *B. distachyon* and *S. moellendorffii*) or PAM fluorometry (AquaPen AP110/C, Photon Systems Instruments) (OD680, maximum fluorescence quantum yield (Qy_max) for *K. nitens*).

Three biological replicates per stress condition were harvested and flash-frozen in liquid nitrogen. Total RNA was extracted using the Spectrum Plant Total RNA Kit (Sigma-Aldrich) with on-column DNase digestion. Libraries were prepared using the Fast RNA-seq Lib Prep Kit V2 (Abclonal) with polyA capture (VAHTS mRNA Capture Beads) and sequenced on the Illumina NovaSeq 6000 (paired-end 150 bp, 20 million reads per sample) at NovogeneAIT Genomics, Singapore.

### Marker gene validation, orthogroup construction and differential gene expression analysis (Figure 1)

Forty canonical Arabidopsis stress marker genes were compiled from the literature, covering 10 stress categories. Each candidate was verified against TAIR GO annotations (ATH_GO_GOSLIM.txt), retaining only genes with experimental evidence codes (IMP, IDA, IEP, or IGI) for a GO term matching the assigned stress category. Orthologs were identified using OrthoFinder v 2.5.5 (Emms & Kelly, 2019), resulting in orthogroups (275,222 groups). For each verified marker, the orthogroup containing its Arabidopsis locus ID was identified, and all member gene IDs were extracted. For each species, stress, and experiment, the ortholog with the largest absolute log2FC was retained when multiple orthologs existed within the same orthogroup.

### Conservation score and classification (Figure 2)

For each orthogroup, a conservation score was computed as the number of species containing at least one DEG member divided by 36 (total species in the atlas). Orthogroups were classified as universal (conservation score > 0.5, corresponding to > 18 species), moderate (0.1 to 0.5), or lineage-specific (< 0.1, corresponding to fewer than 4 species). Per-stress conservation profiles were computed by applying the same classification independently for each stress type.

### Phylostratigraphic analysis (Figure 2)

Each orthogroup was assigned an evolutionary age (phylostratum) based on the most basal taxonomic group represented among its member species. Species were classified into seven major clades (Chlorophyte, Charophyte, Bryophyte, Lycophyte, Gymnosperm, Monocot, Eudicot) following APG IV relationships. Phylostrata ranged from PS1 (Viridiplantae, containing Chlorophyte members) to PS7 (class-specific, restricted to monocots or eudicots). Response strength was measured as the median absolute log2 fold change across all DEGs per orthogroup. Associations between phylostratum rank and response metrics were tested using Spearman rank correlation and Kruskal-Wallis tests.

### Cross-organ and organ-restricted overlap coefficient distributions (Figure 3)

Overlap coefficients were calculated for up-regulated and down-regulated DEGs for all unique control/treatment pairs within organs (organ-restricted) or between organs, excluding identical organs (cross-organ), for each stress type. The shuffled distributions adhere to the same conditions, but also allow for comparisons to be made across stresses. Monte Carlo analysis was performed by comparison against a null distribution generated by shuffling the overlap coefficients over 1000 iterations.

### Clade-specific overlap coefficient distributions (Figure 3)

1-to-1 mappings of every gene in every species were obtained against every species using 1-to-1 orthologues obtained from OrthoFinder. For the entire dataset of DEG lists, a species-specific dictionary was constructed which contained the corresponding 1-to-1 orthologues of that particular species. Next, within each species-specific dictionary, OCs were obtained for every species with every other species, without any organ restrictions, but only allowing cross-clade OCs to be obtained. This allows for basal clades without organs such as leaves and roots to be compared to more modern clades, but again also importantly prevents clade bias as OCs in large clades such as Monocot and Dicot are not allowed to be obtained from within themselves. This process was repeated for each species-specific dictionary, and the final distribution agglomerated in a stress-specific manner and plotted for up- and down-regulated DEGs separately. The mean OCs were then computed and tested for significance with a cross-stress distribution treated in the same way.

### Stress response conservation decays (Figure 3)

Pairwise divergence times between the 36 species were estimated from TimeTree ((Kumar et al., 2022); timetree.org) by assigning each species to a resolved taxonomic tree and using the median estimated age of the most recent common ancestor (MRCA) for each pair. To assess the relationship between evolutionary distance and stress response conservation, Spearman rank correlations were computed between divergence time and mean OC for each stress and direction. Species pairs were binned by divergence time (<10, 10-30, 30-60, 60-100, 100-150, 150-350, 350-500, 500-700, >700 Mya) and mean OC with standard error was computed per bin across all stresses.

### Pathway enrichment analysis (Figure 4)

MapMan bin annotations were assigned to all genes across 36 species using Mercator4 v7.0 (Schwacke et al., 2019). Pathway enrichment analysis was conducted on the Mapman bin assignments in the combined up- and down-regulated DEG list for each experiment using a hypergeometric test against their respective species-specific Mercator bin distribution. FDR-corrected scores for bins containing significantly up (US), down (DS), up-and-down (UDS) and non-significant (NS) regulated bins were assigned per experiment. For each unique species, stress, and organ combination (hereafter “class”), counts of US, DS, UDS or NS were obtained for each bin and normalised by the number of experiments in each class. This step normalises for sample number bias in species, stresses or organs. UP (US+UDS) and DOWN (DS+UDS) scores were obtained and used for downstream analysis.

### MapMan bin directional bias (Figure 4)

For each MapMan bin and stress and/or organ (Level 1), a direction bias was calculated as the difference between the pathway enrichment UP score and DOWN score, and the mean across classes calculated. Positive bias indicates that the bin more frequently contains upregulated DEGs than downregulated DEGs across experiments.

### Functional pathway co-occurrence network construction (Figure 4)

For each class, a binary co-occurrence matrix was constructed: for every pair of MapMan bins (i, j), the matrix entry was set to 1 if both bins contained UP or DOWN scores > 0, and 0 otherwise. Separate matrices were built for UP and DOWN scores. These binary matrices were summed across all classes within a stress and divided by the total number of classes to yield a co-occurrence frequency ranging from 0 to 1. A co-occurrence frequency of 0.30 for a given bin pair under heat stress indicates that in 30% of heat experiments, both bins were simultaneously up or down-regulated. For the normalised all-stress networks, per-stress co-occurrence matrices were first computed independently, then averaged across stresses to prevent stresses with more classes from dominating the signal.

### Universal and stress-specific edge identification (Figure 4)

Universal edges were defined as bin pairs classified as UP in all six stresses or DOWN in all six stresses. Stress-unique edges were defined as edges classified as UP (or DOWN) in exactly one stress and not in any other.

### Hormone functional co-occurrence network analysis (Figure 4)

Phytohormone action first-neighbour subnetworks (Level 1: abscisic acid, salicylic acid, brassinosteroid, gibberellin, jasmonic acid, ethylene, auxin, cytokinin) were extracted from Mercator co-occurrence networks. For each stress, edges were filtered to the top 0.5% by weight, computed separately for UP- and DOWN-regulated edges (99.5th percentile threshold). UP versus DOWN edge weights were used to construct the pie chart. Numbers below each pie indicate the summed UP (left) and DOWN (right) edge weights above threshold. Grey circles indicate hormones present in the network but with no edges above threshold in that stress.

### Knowledge graph validation (Figure 4)

Hormone-stress associations were cross-referenced with the PlantConnectome 2025 knowledge graph (4,819,240 edges extracted from published literature) (S. C. Lim et al., 2025). For each hormone, KG entries were identified by text matching on resolved entity names (e.g., “abscisic acid”, “salicyl*”, “brassinosteroid”). Stress-specific KG entries were identified by co-occurrence of hormone and stress terms (e.g., “heat”, “thermotolerance”, “drought”, “pathogen”, “defense”) within the same KG edge. Gene-level validation was performed by extracting Arabidopsis AGI identifiers from KG gene alias fields and mapping them to MapMan bins via Mercator4 annotations, enabling comparison of KG-predicted hormone-pathway associations with transcriptomically observed co-occurrence patterns.

### Phylogenetic conservation analysis (Figure 4)

Pathway enrichment analysis was performed for seven plant clades (Chlorophyte, Charophyte, Bryophyte, Lycophyte, Gymnosperm, Dicot, Monocot) using the normalised clade-specific data. For each MapMan bin in each class per clade (Level 1), a direction bias was calculated as the difference between the pathway enrichment UP score and DOWN score, and the mean across classes calculated.

### Co-expression network construction and stress module detection (Figure 5)

Gene co-expression networks were constructed for 36 plant species using TEA-GCN (P. K. Lim et al., 2026), which computes z-score standardized mutual rank co-expression scores from TPM-normalized RNA-seq expression matrices. For each species and stress combination, the network was filtered to edges with z >= 1.7 between differentially expressed genes, and Louvain community detection (resolution = 1.0) was applied to identify stress-responsive co-expression modules.

### Cross-species module conservation and null model (Figure 5)

Genes were mapped to 275,222 orthogroups (OrthoFinder). Module pairs from different species under the same stress were compared using the overlap coefficient:

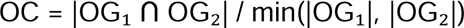

Pairs with OC >= 0.5 and >= 10 shared OGs were considered conserved. To test significance, gene-to-OG assignments were shuffled within each species (100 permutations) and overlap coefficient and direction concordance rates were recomputed. Observed values were compared to the null using Wilcoxon signed-rank tests.

### Within-species module duplication and regulatory divergence (Figure 5)

All module pairs within the same species and stress were compared using the overlap coefficient. Pairs with OC >= 0.3 were classified as duplicated. For each shared OG, DEG directions of all gene copies in each module were compared to compute concordance (same majority direction) and divergence rates. Shared OGs were visualized as paired pie charts scaled by gene copy number with slices reflecting up/down proportions across all copies.

### WGD ohnolog analysis (Figure 5)

Intra-species syntenic anchor pairs were obtained from PLAZA dicots 5.0 and monocots 5.0 for 10 species with matching gene identifiers. Each ohnolog pair was classified by whether both genes co-localize in the same module, split across a duplicated module pair, or split across unrelated modules. A null distribution was generated by shuffling ohnolog pairings within each species (100 permutations), preserving the WGD gene set but randomizing pair assignments.

### Gene sequence extraction and labeling (Figure 6)

Genomic sequences spanning 5 kb upstream and 5 kb downstream of the transcription start site (TSS) were extracted for all protein-coding genes from 25 plant species using Ensembl Plants (release 57) and NCBI Datasets for species lacking Ensembl annotations. For large genomes (*Picea abies, Vitis vinifera*), sequences were extracted directly from chromosome-level assemblies using gene coordinates from GFF3/GTF annotations. For each stress and regulation direction, genes were labeled as stress-responsive (top quartile of DEG frequency across experiments) or non-responsive (bottom quartile). When quartile-based labeling collapsed to fewer than two classes, a binary median split was applied.

### Orthogroup-based data splitting (Figure 6)

To prevent information leakage from homologous genes, all train/validation/test splits were performed at the orthogroup level using OrthoFinder-derived orthogroups (2.5M gene-to-orthogroup mappings). Orthogroups were randomly assigned to train (70%), validation (15%), or test (15%) partitions, ensuring that all genes within an orthogroup were assigned to the same partition.

### CNN baseline (Figure 6)

A convolutional neural network with three Conv1D blocks (64, 128, 64 filters; kernel size 8; ReLU activation; max pooling) followed by flattening, a 128-unit dense layer, a 64-unit dense layer, and a sigmoid output was trained on 600 bp one-hot encoded DNA sequences centered on the TSS (positions 4700-5300 in the 10 kb FASTA). Training used binary cross-entropy loss, Adam optimizer (learning rate 1e-3), batch size 128, and early stopping on validation auROC with patience of 10 epochs. Training data were balanced by undersampling the majority class.

### PlantCAD2 (DNA foundation model) (Figure 6)

PlantCAD2 is a bidirectional Mamba/Caduceus architecture (88M parameters) pre-trained on 65 plant genomes. The pre-trained model was obtained from the authors’ repository. For fine-tuning, 600 bp promoter DNA sequences were tokenized using the PlantCAD2 tokenizer. Low-rank adaptation (LoRA) with rank r=8 was applied to the model’s linear layers, yielding 2.4M trainable parameters. A classification head consisting of mean pooling followed by a two-layer MLP (128 units, ReLU, dropout 0.1, sigmoid output) was appended. Training used binary cross-entropy loss, AdamW optimizer (learning rate 2e-5, weight decay 0.01), batch size 16, and early stopping on validation auROC. Models were trained on Google Colab (NVIDIA A100 40GB) with mixed precision (bfloat16). Only input_ids were passed to the model (no attention_mask or token_type_ids, as required by the Caduceus architecture).

### PlantRNA-FM (RNA foundation model) (Figure 6)

The pre-trained model was loaded from the HuggingFace repository (yangheng/PlantRNA-FM). For fine-tuning, 1,024 bp mRNA sequences starting from the TSS (positions 5000-6024 in the 10 kb FASTA) were tokenized using the PlantRNA-FM tokenizer. LoRA with rank r=16 was applied, yielding 369K trainable parameters. A classification head (mean pooling, 128-unit dense, ReLU, dropout 0.1, sigmoid) was appended. Training used binary cross-entropy loss, AdamW optimizer (learning rate 2e-5), batch size 16, and early stopping. Model weights were loaded with strict=False to accommodate extra position embedding keys.

### Late fusion (Figure 6)

DNA and RNA model predictions were fused. Two fusion strategies were evaluated: (1) simple averaging of predicted probabilities, and (2) logistic regression on the two prediction scores trained on the validation set. Both strategies yielded near-identical performance (data not shown); simple averaging was used for all reported results.

### Transcript window scan (Figure 6)

To identify which transcript regions carry regulatory information, PlantRNA-FM was retrained independently on four 1 kb windows relative to the TSS: 5’ end (0 to +1 kb), mid-transcript (+0.5 to +1.5 kb), deep transcript (+1 to +2 kb), and 3’ end (last 1 kb before transcription termination site). For each window, the sequences were tokenized, and used to train and evaluate a fresh PlantRNA-FM model with the same architecture and hyperparameters.

### Interpretable feature-based models (Figure 6)

Three groups of interpretable features were computed for each gene: (1) Dinucleotide frequencies: all 16 dinucleotide frequencies in both promoter (600 bp) and mRNA (1,024 bp) regions, plus GC content (34 features total). (2) Motif counts: occurrences of 16 known plant cis-regulatory elements (ABRE, DRE/CRT, W-box, GCC-box, G-box, TATA-box, CAAT-box, MBS, as-1, MYC, LTR, ARE, AU-rich element, PUF-binding site, DST element, poly(A) signal) plus the top 10 discovered k-mers (k=8) from each model’s attribution analysis (CNN, PlantCAD2, PlantRNA-FM), counted in both promoter and mRNA regions with reverse complement matching (∼92 features after deduplication). (3) RNA secondary structure: minimum free energy (MFE) per nucleotide, fraction of paired bases, number of stems and hairpins, maximum stem length, G-quadruplex potential, C-rich runs, and AU-rich stretches, computed on the first 200 bp of mRNA using ViennaRNA (8 features). Structure features were precomputed and cached for all ∼970,000 genes in the dataset.

Gradient-boosted machines (sklearn GradientBoostingClassifier; 300 estimators, max depth 5, learning rate 0.1, min samples leaf 10, subsample 0.8) were trained on each feature group individually and on all features combined. Training data were balanced by undersampling, and features were standardized (zero mean, unit variance).

### Model interpretability (Figure 6)

CNN attributions were computed using integrated gradients with 50 interpolation steps between a zero baseline and the input one-hot encoding. PlantCAD2 attributions were obtained via in silico mutagenesis (ISM), systematically mutating each position and recording the change in predicted probability. PlantRNA-FM attributions were computed using gradient x input. For each model, the top 200 highest-attribution sequences per stress per direction were extracted and scanned for matches to known cis-regulatory elements (exact substring match including reverse complement).

### Statistical analysis of sequence features (Figure 6)

For each of 40 sequence features (dinucleotide frequencies, GC content, motif counts, RNA structure descriptors), Mann-Whitney U tests were performed comparing feature values between stress-responsive and non-responsive genes (n=10,000 per group, randomly sampled). Tests were conducted separately for each stress, direction, and feature combination (400 tests total). P-values were corrected using the Bonferroni method. Effect sizes were quantified using Cohen’s d. Feature importance was assessed using mean Gini importance from the combined GBM model, averaged across all stress-direction combinations.

### Software and visualization

All analyses were performed in Python 3 using pandas, NumPy, SciPy, matplotlib, and seaborn.

## Supplementary tables

**Table S1**. **Curated metadata for stress transcriptome experiments.** Complete sample-level metadata for all RNA-seq experiments included in the atlas, listing SRA run accession, species, BioProject, stress type, sample organ, differentiating conditions between control and treatment samples, experiment group assignment, and control/treatment designation. Each row represents one sample. Experiment groups define sets of samples sharing a common experimental design within a BioProject; control/treatment pairs within each group were manually curated by inspecting sample descriptions and experimental conditions.

**Table S2**. **GO-verified canonical stress marker genes used for pipeline validation.** Arabidopsis marker genes across stress categories used to validate the cross-species DEG calling pipeline (Fig. 1D). Marker genes were selected from curated literature candidates and retained only if annotated with experimental GO evidence codes (IMP, inferred from mutant phenotype; IDA, inferred from direct assay; IEP, inferred from expression pattern; IGI, inferred from genetic interaction) for a GO term matching the assigned stress category in the TAIR ATH_GO_GOSLIM annotation file. Columns: stress category, gene name, AGI locus identifier, matching GO term(s), and supporting evidence code(s).

**Table S3. Orthogroup stress response matrix.** Each row represents one of 101,692 stress-responsive orthogroups. Columns correspond to the 36 species in the atlas. Cell values list the stress types (comma-separated) for which at least one gene in that orthogroup was differentially expressed (|log2FC| > 1, adjusted p < 0.05) in that species. Empty cells indicate no significant differential expression was detected. Summary columns: n_species, number of species with at least one DEG; n_stresses, number of distinct stress types across all species. Orthogroups are sorted by decreasing species breadth.

**Table S4. Orthogroup phylostratum assignments and taxonomic distribution.** Phylostratum classification for each orthogroup, assigned based on the most basal taxonomic group containing members of the orthogroup. Columns: orthogroup identifier, phylostratum, number of species represented in the orthogroup, number of clades represented, and list of clades containing member genes.

**Table S5. Direction bias of MapMan functional categories across stress types.** Each row represents a MapMan Level 1 bin under a given stress condition. Columns: PARENT_BINCODE, MapMan bin code; bin_name, full bin path; short_name, abbreviated bin name; stress, stress type (Cold, Drought, Heat, Heavy metal, Pathogen, or Salt); mean_UP and mean_DOWN, mean fraction of genes up- or down-regulated across experiments for that bin-stress combination; mean_bias, directional bias (mean_UP minus mean_DOWN); n_experiments and n_species, number of experiments and species contributing to that bin-stress estimate; overall_mean_UP, overall_mean_DOWN, and overall_mean_bias, corresponding values averaged across all six stress types.

**Table S6. Stress-specific co-occurrence network edges between MapMan functional categories.** Each row represents an edge linking two Mercator functional annotation bins that are frequently co-regulated under a given stress. source and target, the two MapMan bins connected by the edge (format: Parent category.subcategory); weight, co-occurrence frequency of genes between the two bins across all organs; Master bincode, Mercator hierarchical bin code of the source node; direction, regulation direction of the underlying genes (UP or DOWN); stress, stress condition under which the network was constructed (Cold, Drought, Heat, Heavy metal, Pathogen, or Salt).

**Table S7. Cis-regulatory motifs discovered by attribution analysis.** Top 10 most frequent 8-mers extracted from the 200 highest-attribution sequences per model, stress, and regulation direction. Attributions were computed using integrated gradients (CNN), in silico mutagenesis (PlantCAD2), and gradient x input (PlantRNA-FM). Canonical form is the lexicographically smaller of the k-mer and its reverse complement, used for deduplication. Rank indicates frequency rank within each model-stress-direction combination.

**Table S8. Cohen’s d effect sizes for genomic features of stress-responsive genes.** Each row represents one stress type, regulation direction (UP/DOWN), and genomic feature combination. resp_mean and nonr_mean: mean feature values for responsive and non-responsive genes, respectively (n = 10,000 each, subsampled). direction_vs_nonresp: whether the feature is higher or lower in responsive genes. cohens_d: Cohen’s d effect size. effect_size_rb: rank-biserial effect size. p_value: Wilcoxon rank-sum test p-value; p_corrected: Bonferroni adjusted p-value. Significance: * p < 0.05, ** p < 0.01, *** p < 0.001.

**Table S9. Predictive feature importances for stress-responsive gene classification.** Each row represents one genomic feature, stress type, and regulation direction (UP/DOWN). importance: feature importance score from a trained classifier predicting stress-responsive vs non-responsive genes. Features are derived from mRNA and promoter sequence composition (e.g., dinucleotide frequencies). Higher values indicate greater contribution to distinguishing stress-responsive genes from non-responsive genes.

## Supplementary Figures

**Figure S1. Phenotypic measurements of stress treatments in *Brachypodium distachyon*.** Plants were subjected to cold, drought, heat and salt stresses as described in the Methods. Plants were photographed and various phenotypic characteristics such as max shoot length, max root length, surface area and fresh weight were measured at 11 and 17 DAG (1 and 7 days after treatment).

**Figure S2. Phenotypic measurements of stress treatments in *Selaginella moellendorffii*.** *Selaginella moellendorffii* plants were subjected to cold, drought, heat and salt stresses as described in the Methods. Plants were photographed and average leaf surface area was measured (Methods) at 9 and 16 DAS (1 and 7 days after treatment).

**Figure S3. Phenotypic measurements of stress treatments in *Klebsormidium nitens*.** *Klebsormidium nitens* cultures were subjected to cold, drought, heat and salt stresses as described in the Methods. Growth density (OD 680) and maximum fluorescence quantum yield (Qy_max) measured were obtained (Methods) at 0, 3, 4, 6 days after treatment.

**Figure S4. Pairwise overlap coefficients of stress-responsive one-to-one orthologues as a function of species divergence time (Mya) for six stresses (panels).** Each point represents the mean OC for one species pair under one stress and direction (UP-regulated, red; DOWN-regulated, blue). LOWESS curves (fraction = 0.4) are overlaid to visualize the trend. Spearman rank correlation coefficients (rho) and significance levels are shown per panel. All UP-regulated correlations are significantly negative (p < 0.001), indicating phylogenetic decay of stress response conservation. DOWN-regulated genes show weaker decay, with heat showing no significant correlation (rho = -0.08, ns). Pathogen UP-regulated genes show the steepest decay (rho = -0.43), consistent with lineage-specific defense activation. Divergence times were estimated from TimeTree (Kumar et al. 2022). Only public and in-house data for the six most broadly sampled stresses (>=22 species each) are shown.

**Figure S5. Phylogenetic conservation of directional bias.** Phylogenetic conservation of direction bias for key MapMan Level 1 bins across seven plant clades arranged from basal (left) to derived (right): Chlorophyte (1 species), Charophyte (1 species), Bryophyte (2 species), Lycophyte (1 species), Gymnosperm (1 species), Dicot (18 species), Monocot (12 species). Values represent the summed direction bias (UP - DOWN scores) for each bin, averaged across all stresses within each clade. Red indicates UP-biased (more frequently co-upregulated); blue indicates DOWN-biased (more frequently co-downregulated). Protein quality control is UP-biased across all clades including Chlorophyte, indicating deep conservation of the chaperone stress response.

**Figure S6. Stress-specific functional co-regulation networks for cold, drought, heavy metal, pathogen and salt.** Nodes represent Mercator Level 1 functional bin categories; node size is proportional to total weighted degree within that stress’s unique network. Edge colour indicates direction of co-regulation: red = co-upregulated, blue = co-downregulated; edge width and colour intensity are proportional to edge weight.

**Figure S7. Knowledge graph validation of hormone-stress associations.** Left: number of literature entries linking each hormone to each stress in the PlantConnectome knowledge graph (4.8M edges, predominantly Arabidopsis-derived). Right: transcriptomic direction bias from our cross-kingdom dataset. Comparison reveals literature bias toward pathogen-hormone interactions and identifies broadly responsive hormones (SA, BR) underrepresented in current literature.

**Figure S8. Z-score distributions of TEA-GCN co-expression networks across 36 plant species.** Each panel shows the distribution of z-score standardized mutual rank co-expression scores for edges passing the initial threshold (z >= 1.5). Dashed black line indicates the z = 1.7 threshold used for stress module detection. Red dotted line marks the maximum z-score per species. Five species (red histograms, bold names) were excluded from downstream module analyses because their z-score distributions did not reach the 1.7 threshold: Brassica oleracea (max z = 1.61), Lactuca sativa (max z = 1.70), Medicago truncatula (max z = 1.68), Physcomitrium patens (max z = 1.63), and Spinacia oleracea (max z = 1.57). This truncation likely reflects lower RNA-seq sample counts in the TEA-GCN input, which compresses the mutual rank z-score distribution. The remaining 31 species were used for all module-level analyses.

**Figure S9. Distribution of overlap coefficient (top) and concordance rate (bottom) across all module pairs for cross-species (blue) and within-species (red) comparisons.** Dark shades show observed values; light shades show the null distribution from shuffled gene-to-OG assignments (100 permutations). Dashed and dotted vertical lines indicate means.

## Supplementary Datasets

**Supplementary Dataset 1. OrthoFinder orthogroup assignments.** Orthogroup membership for all genes across all species in the atlas, as computed by OrthoFinder. Each line contains an orthogroup identifier followed by the gene IDs of all member genes across species, with transcript-level suffixes retained from the original genome annotations.

**Supplementary Dataset 2. Kingdom-wide differentially expressed genes.** DESeq2 results for all control/treatment comparisons across species and stress conditions. Columns: species, stress type, BioProject accession (or in-house experiment identifier), sample organ, experiment group, experiment identifier, direction of regulation, gene identifier, log2 fold change, raw p-value, adjusted p-value, and whether the data originated from in-house experiments.

**Supplemental Dataset 3. Thresholded co-expression networks for 36 plant species.** Archive contains one tab-delimited file per species ({Species}_edges.tsv), each listing all gene pairs with a TEA-GCN z-score standardized mutual rank co-expression score above 1.5. Columns: source (gene ID), target (gene ID), z-score of the mutual rank co-expression strength.

**Supplementary Dataset 4. Cross-species co-expression modules and conservation analysis.** Archive containing two files: (1) gene-to-module assignments for each species, listing every gene and its co-expression module membership; and (2) pairwise edges between modules across species, with overlap coefficients computed on shared orthogroup content. Conserved module pairs (overlap coefficient >= 0.5) were identified and clustered using Louvain community detection to define groups of functionally related modules across species.

**Supplementary Dataset 5. Trained model weights for stress-responsive gene prediction.** Archive containing fine-tuned model weights for three architectures trained to predict stress-responsive genes from DNA or RNA sequence across 25 plant species. (a) cnn_models/: 10 CNN baseline models (Keras format, 3x Conv1D on 600 bp one-hot DNA, ∼4 MB each). (b) plantcad2_adapters/: 10 LoRA adapter weights for PlantCAD2 (Mamba/Caduceus DNA language model, 88M parameters, r=8, HuggingFace PEFT format, ∼9 MB each). Requires base model kuleshov-group/PlantCaduceus. (c) plantrna_fm_models/: 10 PlantRNA-FM fine-tuned models (ESM RNA language model, 33M parameters, LoRA r=16, PyTorch state_dict, ∼131 MB each). Requires base model yangheng/PlantRNA-FM. Each folder contains one model per stress (Heat, Cold, Drought, Salt, Pathogen) and regulation direction (UP, DOWN). All models were trained using orthogroup-based splits to prevent homology leakage. See README.md in the archive for loading instructions and dependencies.

**Supplementary Dataset 6. Precomputed sequence features for 1,687,829 genes across 25 plant species.** Feature table containing 658 sequence-derived features for all protein-coding genes with sufficient flanking sequence (at least 5 kb upstream and 1 kb downstream of TSS). Columns: gene_id, species, 16 dinucleotide frequencies in the promoter (600 bp upstream of TSS), 16 dinucleotide frequencies in the mRNA (1,024 bp from TSS), GC content for both regions, counts of 16 known cis-regulatory elements (ABRE, DRE/CRT, W-box, GCC-box, G-box, TATA-box, CAAT-box, MBS, as-1, MYC, LTR, ARE, AU-rich element, PUF-binding site, DST element, poly(A) signal) in both promoter and mRNA regions (32 features), counts of 292 discovered 8-mer motifs from CNN, PlantCAD2, and PlantRNA-FM attribution analysis in both regions (584 features), and 8 RNA secondary structure descriptors (minimum free energy per nucleotide, fraction of paired bases, number of stems, number of hairpins, maximum stem length, G-quadruplex count, C-rich run count, AU-rich stretch count) computed on the first 200 bp of mRNA using ViennaRNA..

## Data availability

All supplemental datasets are available at 10.6084/m9.figshare.31995939, with exception of Supplemental Dataset 3, which is available at: https://sid.erda.dk/share_redirect/GSCXXPyGbM. The scripts made to generate the data and figures are available at https://github.com/mutwil/KingdomStress. The raw sequencing data are available at E-MTAB-16937, E-MTAB-16940, E-MTAB-16941.

