## Supplementary material for "Kingdom-wide comparative transcriptomics reveals deeply conserved and predictable stress response programs across Viridiplantae": Figure S1-9

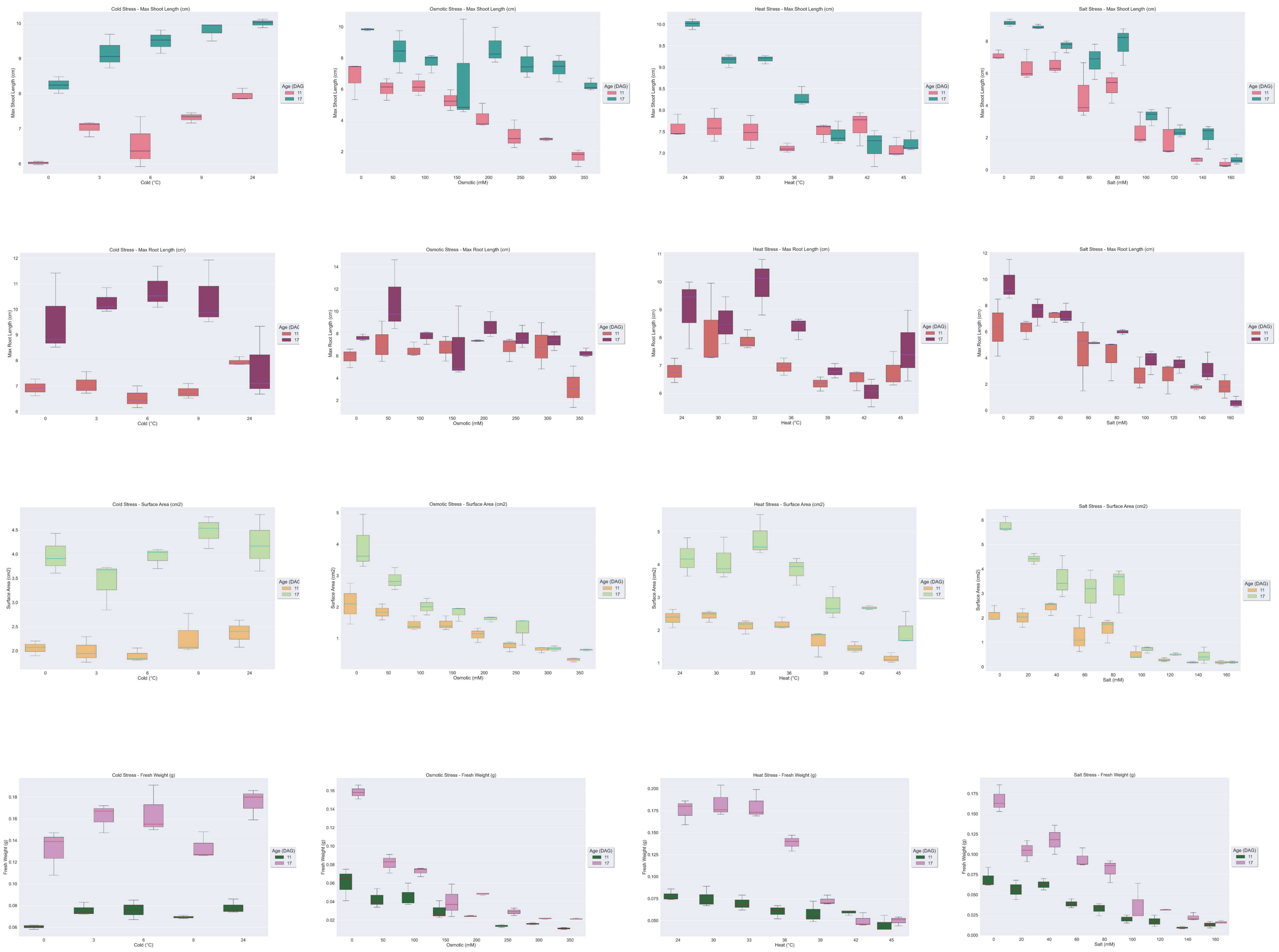
Cold Stress - Max Shoot Length (cm)
Osmotic Stress - Max Shoot Length (cm)
Heat Stress - Max Shoot Length (cm)
Salt Stress - Max Shoot Length (cm)
Cold Stress - Max Root Length (cm)
Osmotic Stress - Max Root Length (cm)
Heat Stress - Max Root Length (cm)
Salt Stress - Max Root Length (cm)
Cold Stress - Surface Area (cm2)
Osmotic Stress - Surface Area (cm2)
Heat Stress - Surface Area (cm2)
Salt Stress - Surface Area (cm2)
Cold Stress - Fresh Weight (g)
Osmotic Stress - Fresh Weight (g)
Heat Stress - Fresh Weight (g)
Salt Stress - Fresh Weight (g)
Max Shoot Length (cm)
Max Root Length (cm)
Surface Area (cm2)
Fresh Weight (g)
Cold (°C)
Osmotic (mM)
Heat (°C)
Salt (mM)
Age (DAG)
11
17

Figure S1

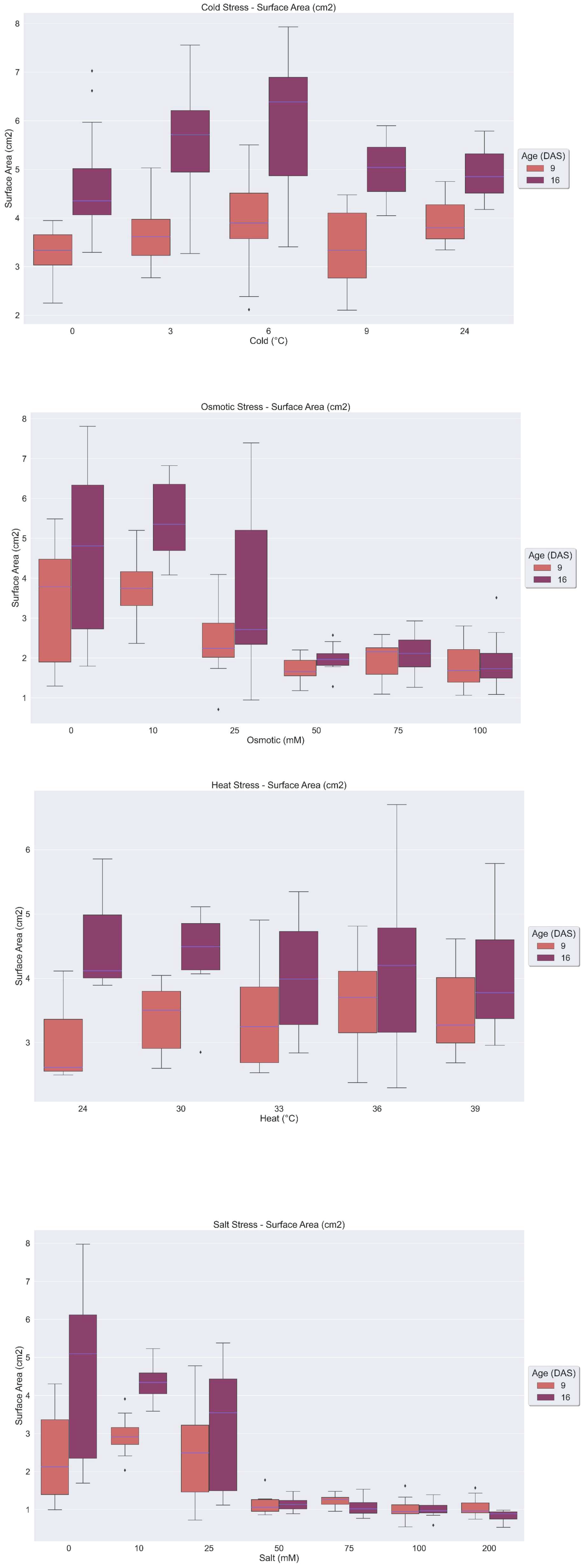
Cold Stress - Surface Area (cm2)
Surface Area (cm2)
Cold (°C)
0
3
6
9
24
Age (DAS)
9
16
Osmotic Stress - Surface Area (cm2)
Osmotic (mM)
0
10
25
50
75
100
Heat Stress - Surface Area (cm2)
Heat (°C)
24
30
33
36
39
Salt Stress - Surface Area (cm2)
Salt (mM)
0
10
25
50
75
100
200

Figure S2

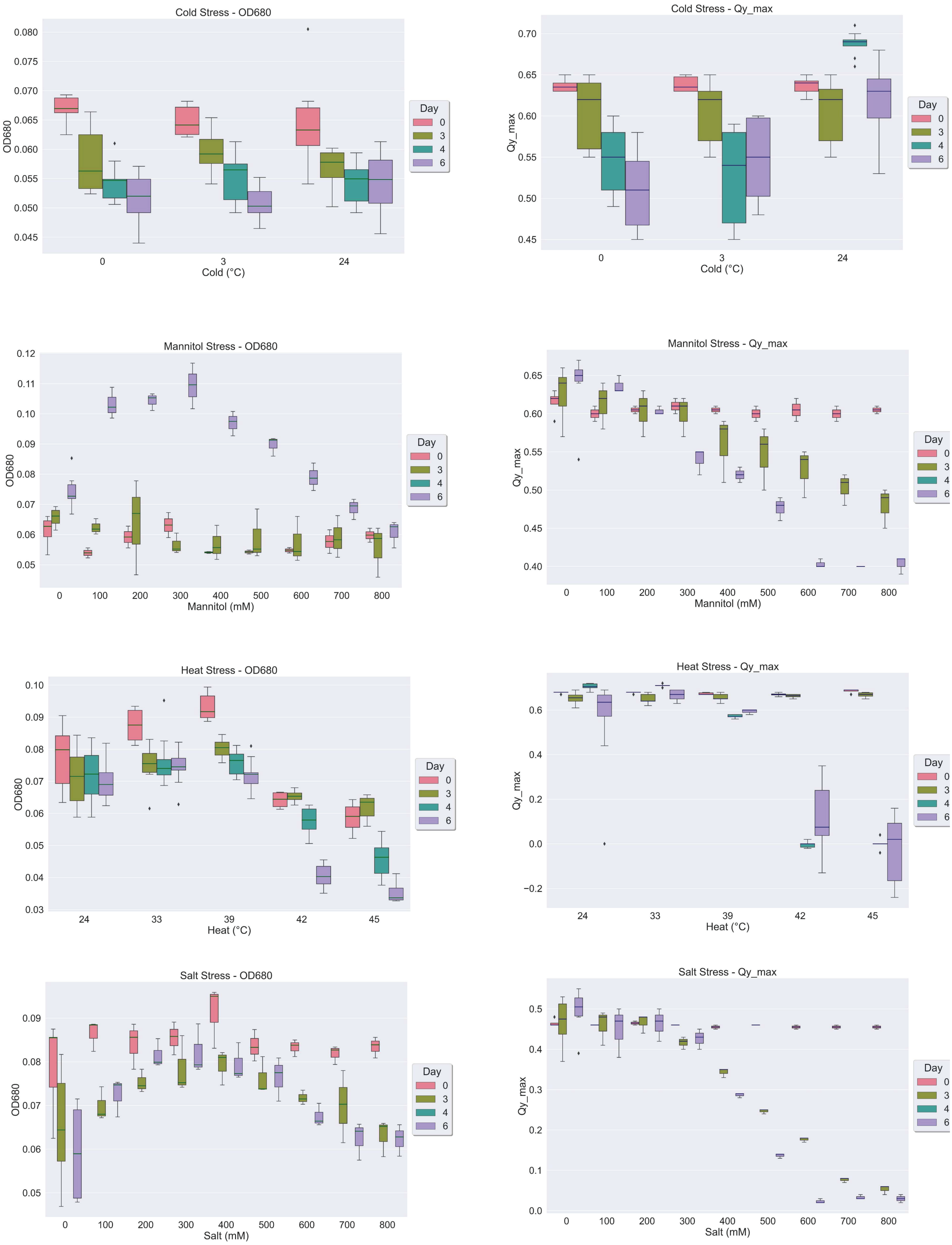
Cold Stress - OD680
OD680
Cold (°C)
0
3
24
Day
0
3
4
6
Cold Stress - Qy_max
Qy_max
Cold (°C)
Mannitol Stress - OD680
OD680
Mannitol (mM)
0
100
200
300
400
500
600
700
800
Mannitol Stress - Qy_max
Qy_max
Mannitol (mM)
Heat Stress - OD680
OD680
Heat (°C)
24
33
39
42
45
Heat Stress - Qy_max
Qy_max
Heat (°C)
Salt Stress - OD680
OD680
Salt (mM)
Salt Stress - Qy_max
Qy_max
Salt (mM)

Figure S3

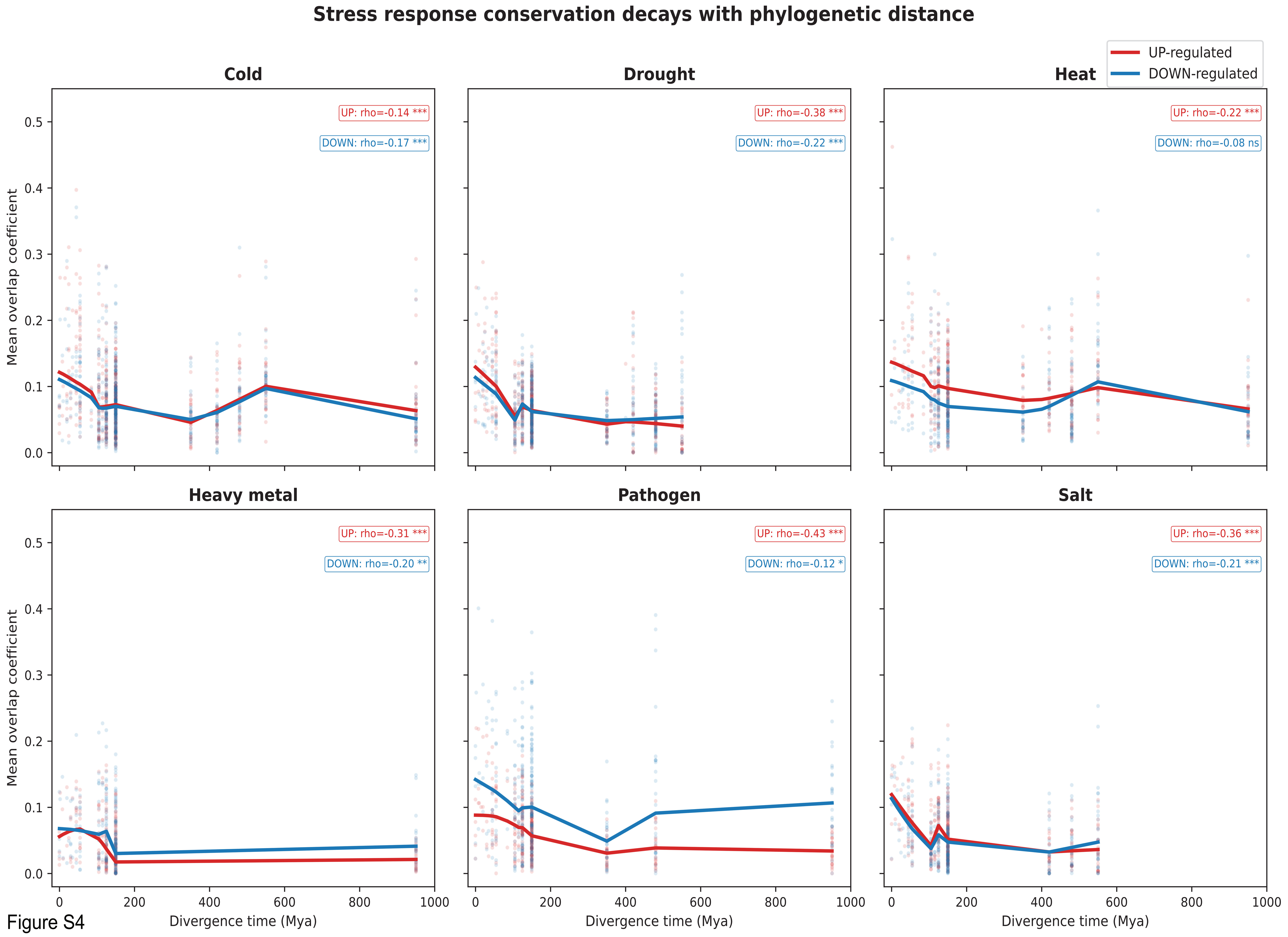
Stress response conservation decays with phylogenetic distance
UP-regulated
DOWN-regulated
Cold
UP: rho=-0.14 ***
DOWN: rho=-0.17 ***
Drought
UP: rho=-0.38 ***
DOWN: rho=-0.22 ***
Heat
UP: rho=-0.22 ***
DOWN: rho=-0.08 ns
Heavy metal
UP: rho=-0.31 ***
DOWN: rho=-0.20 **
Pathogen
UP: rho=-0.43 ***
DOWN: rho=-0.12 *
Salt
UP: rho=-0.36 ***
DOWN: rho=-0.21 ***
Mean overlap coefficient
Divergence time (Mya)

Figure S4

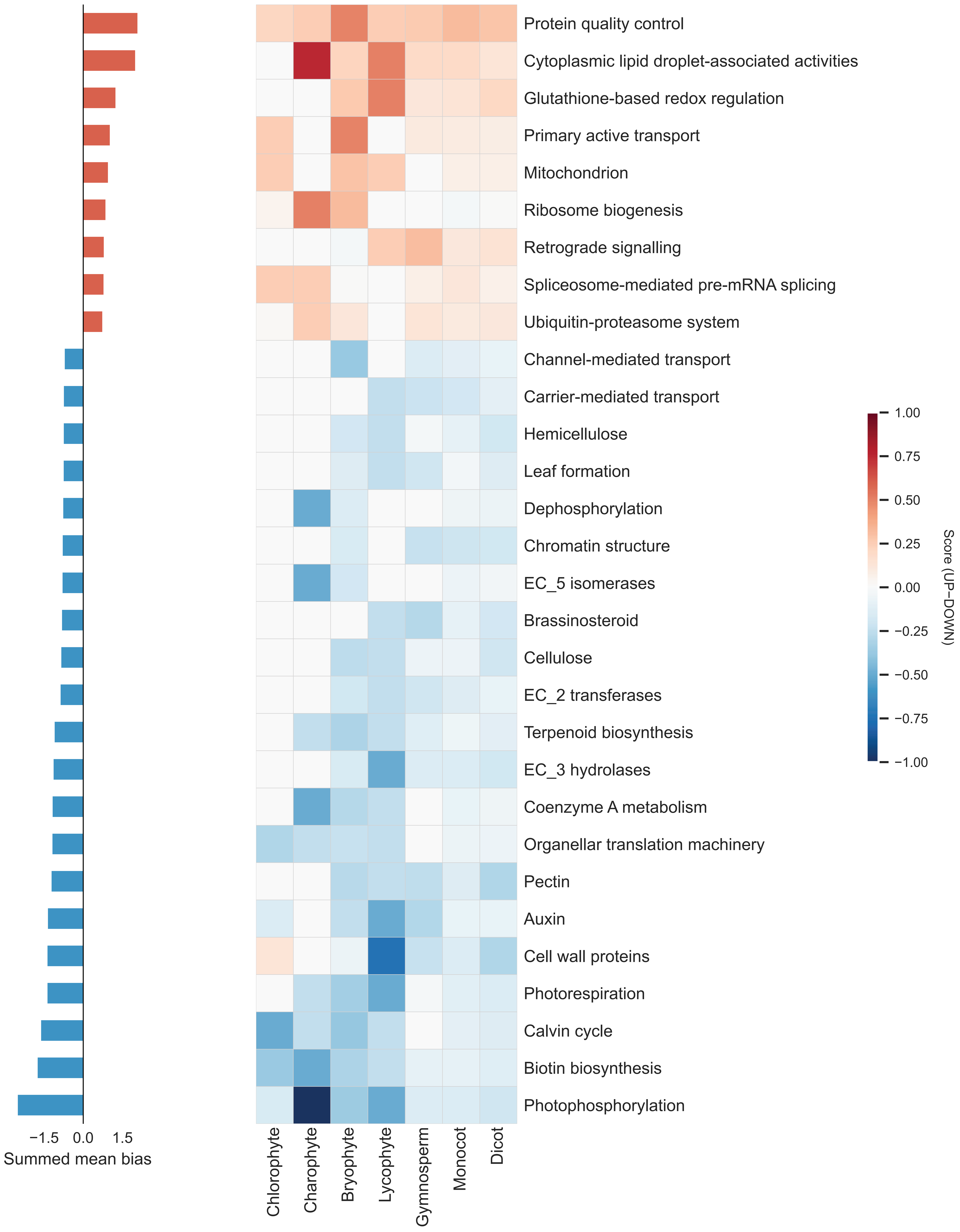
Protein quality control
Cytoplasmic lipid droplet-associated activities
Glutathione-based redox regulation
Primary active transport
Mitochondrion
Ribosome biogenesis
Retrograde signalling
Spliceosome-mediated pre-mRNA splicing
Ubiquitin-proteasome system
Channel-mediated transport
Carrier-mediated transport
Hemicellulose
Leaf formation
Dephosphorylation
Chromatin structure
EC_5 isomerases
Brassinosteroid
Cellulose
EC_2 transferases
Terpenoid biosynthesis
EC_3 hydrolases
Coenzyme A metabolism
Organellar translation machinery
Pectin
Auxin
Cell wall proteins
Photorespiration
Calvin cycle
Biotin biosynthesis
Photophosphorylation
−1.5 0.0 1.5
Summed mean bias
Chlorophyte
Charophyte
Bryophyte
Lycophyte
Gymnosperm
Monocot
Dicot
Clade
1.00
0.75
0.50
0.25
0.00
−0.25
−0.50
−0.75
−1.00
Score (UP−DOWN)

Figure S5

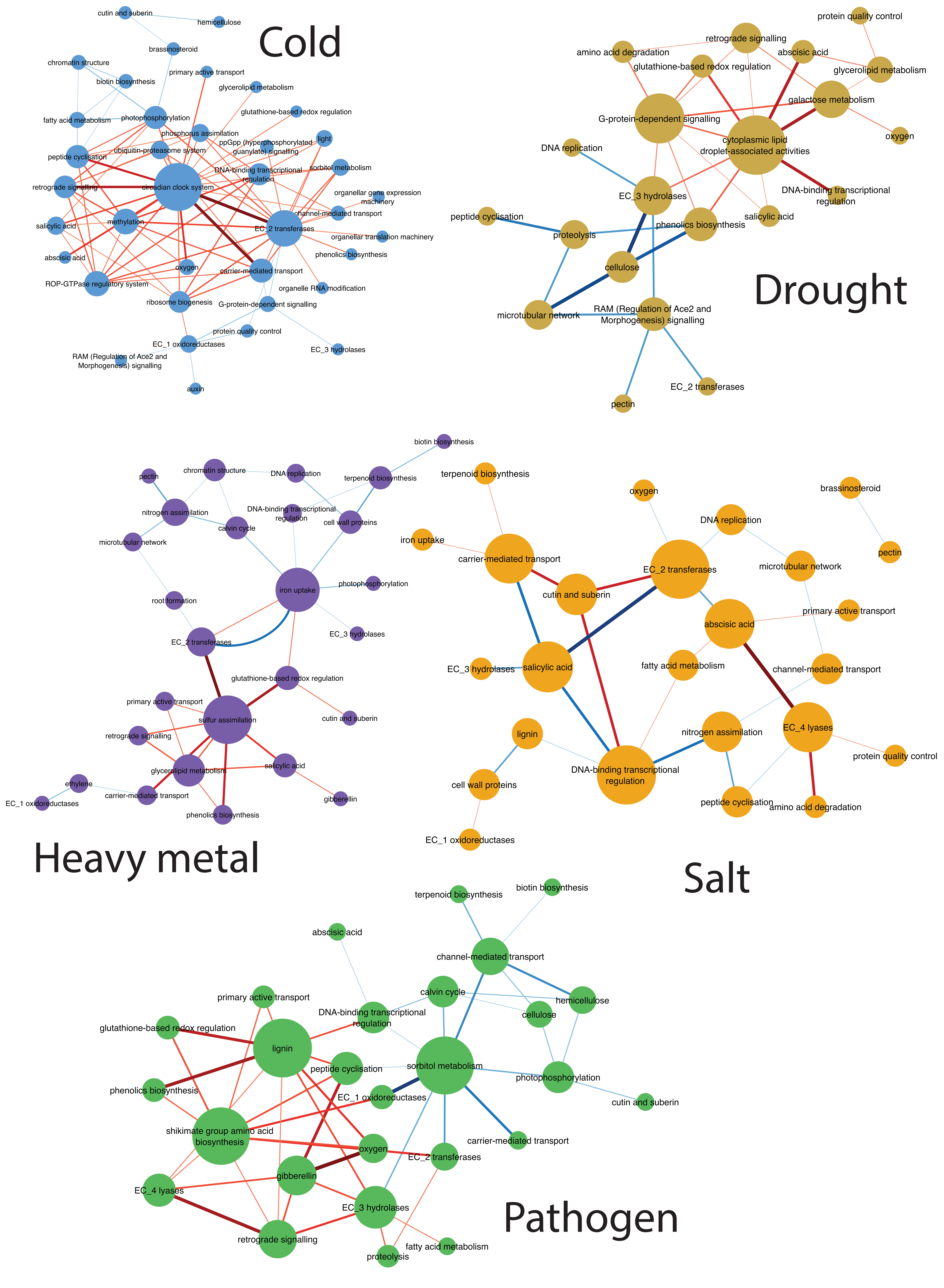

Cold
cutin and suberin
hemicellulose
brassinosteroid
chromatin structure
primary active transport
biotin biosynthesis
glycerolipid metabolism
glutathione-based redox regulation
fatty acid metabolism
photophosphorylation
phosphorus assimilation
light
ppGpp (hyperphosphorylated guanylate) signalling
peptide cyclisation
ubiquitin-proteasome system
DNA-binding transcriptional regulation
sorbitol metabolism
retrograde signalling
circadian clock system
organellar gene expression machinery
channel-mediated transport
salicylic acid
methylation
EC_2 transferases
organellar translation machinery
phenolics biosynthesis
abscisic acid
oxygen
carrier-mediated transport
ROP-GTPase regulatory system
organelle RNA modification
ribosome biogenesis
G-protein-dependent signalling
protein quality control
EC_1 oxidoreductases
EC_3 hydrolases
RAM (Regulation of Ace2 and Morphogenesis) signalling
auxin
Drought
protein quality control
retrograde signalling
amino acid degradation
abscisic acid
glutathione-based redox regulation
glycerolipid metabolism
galactose metabolism
G-protein-dependent signalling
oxygen
DNA replication
cytoplasmic lipid droplet-associated activities
DNA-binding transcriptional regulation
EC_3 hydrolases
peptide cyclisation
salicylic acid
phenolics biosynthesis
proteolysis
cellulose
RAM (Regulation of Ace2 and Morphogenesis) signalling
microtubular network
EC_2 transferases
pectin
Heavy metal
biotin biosynthesis
chromatin structure
DNA replication
pectin
terpenoid biosynthesis
DNA-binding transcriptional regulation
nitrogen assimilation
cell wall proteins
calvin cycle
microtubular network
iron uptake
photophosphorylation
root formation
EC_3 hydrolases
EC_2 transferases
glutathione-based redox regulation
primary active transport
sulfur assimilation
cutin and suberin
retrograde signalling
salicylic acid
ethylene
glycerolipid metabolism
EC_1 oxidoreductases
carrier-mediated transport
gibberellin
phenolics biosynthesis
Salt
terpenoid biosynthesis
oxygen
brassinosteroid
DNA replication
iron uptake
carrier-mediated transport
pectin
EC_2 transferases
microtubular network
cutin and suberin
primary active transport
abscisic acid
EC_3 hydrolases
salicylic acid
fatty acid metabolism
channel-mediated transport
lignin
nitrogen assimilation
EC_4 lyases
protein quality control
DNA-binding transcriptional regulation
cell wall proteins
peptide cyclisation
amino acid degradation
EC_1 oxidoreductases
Pathogen
biotin biosynthesis
terpenoid biosynthesis
abscisic acid
channel-mediated transport
calvin cycle
hemicellulose
primary active transport
DNA-binding transcriptional regulation
cellulose
glutathione-based redox regulation
lignin
sorbitol metabolism
peptide cyclisation
photophosphorylation
phenolics biosynthesis
EC_1 oxidoreductases
cutin and suberin
shikimate group amino acid biosynthesis
carrier-mediated transport
oxygen
EC_2 transferases
gibberellin
EC_4 lyases
EC_3 hydrolases
retrograde signalling
fatty acid metabolism
proteolysis


Figure S6

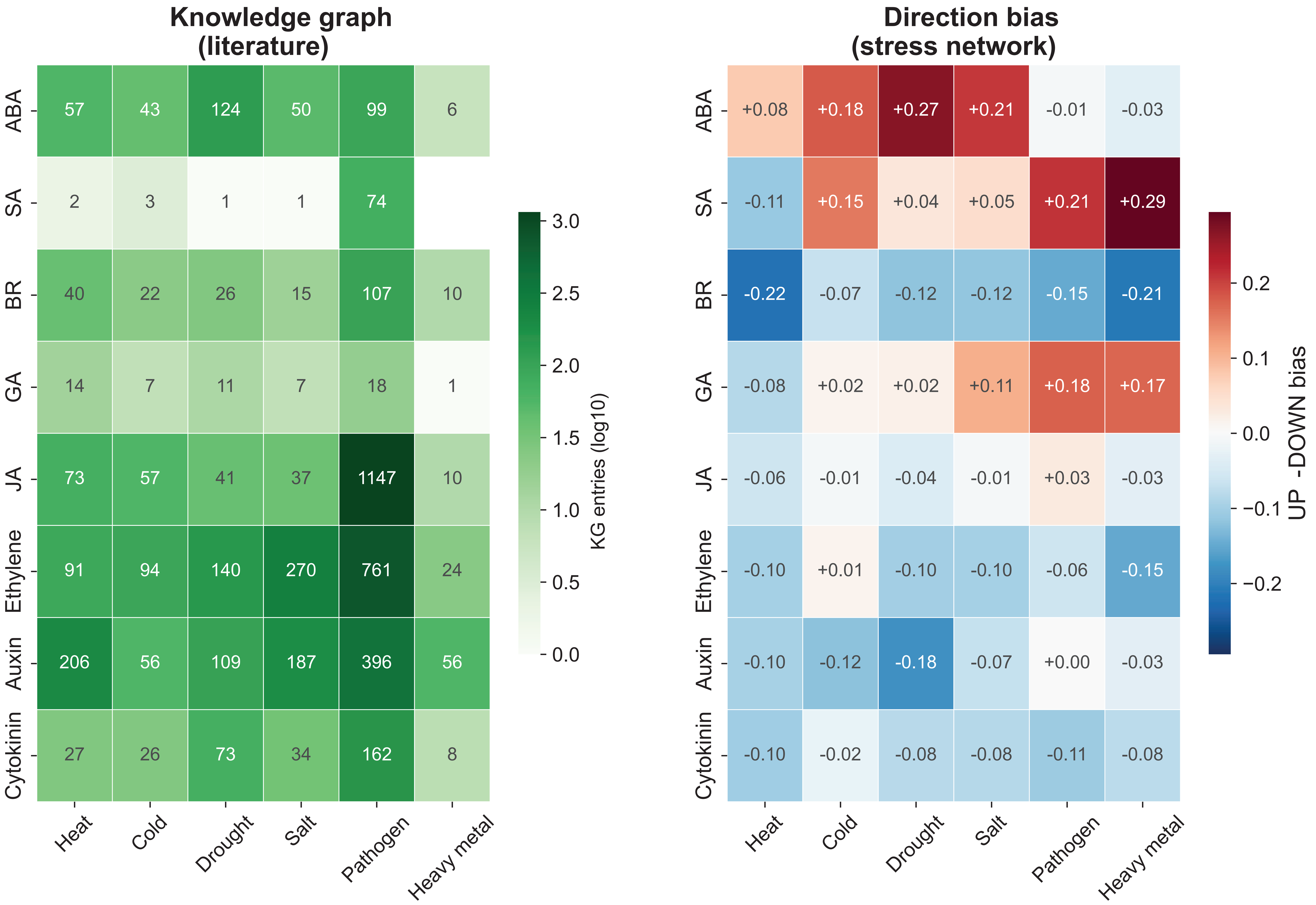
Knowledge graph
(literature)
ABA
SA
BR
GA
JA
Ethylene
Auxin
Cytokinin
57 43 124 50 99 6
2 3 1 1 74
40 22 26 15 107 10
14 7 11 7 18 1
73 57 41 37 1147 10
91 94 140 270 761 24
206 56 109 187 396 56
27 26 73 34 162 8
Heat
Cold
Drought
Salt
Pathogen
Heavy metal
KG entries (log10)
3.0
2.5
2.0
1.5
1.0
0.5
0.0
Direction bias
(stress network)
+0.08 +0.18 +0.27 +0.21 -0.01 -0.03
-0.11 +0.15 +0.04 +0.05 +0.21 +0.29
-0.22 -0.07 -0.12 -0.12 -0.15 -0.21
-0.08 +0.02 +0.02 +0.11 +0.18 +0.17
-0.06 -0.01 -0.04 -0.01 +0.03 -0.03
-0.10 +0.01 -0.10 -0.10 -0.06 -0.15
-0.10 -0.12 -0.18 -0.07 +0.00 -0.03
-0.10 -0.02 -0.08 -0.08 -0.11 -0.08
UP -DOWN bias
0.2
0.1
0.0
−0.1
−0.2

### Figure S7

**Supplementary Figure: Z-score distributions of co-expression networks (z >= 1.5)**
**Dashed line = z = 1.7 module detection threshold. Red species names = excluded (max z < 1.7)**

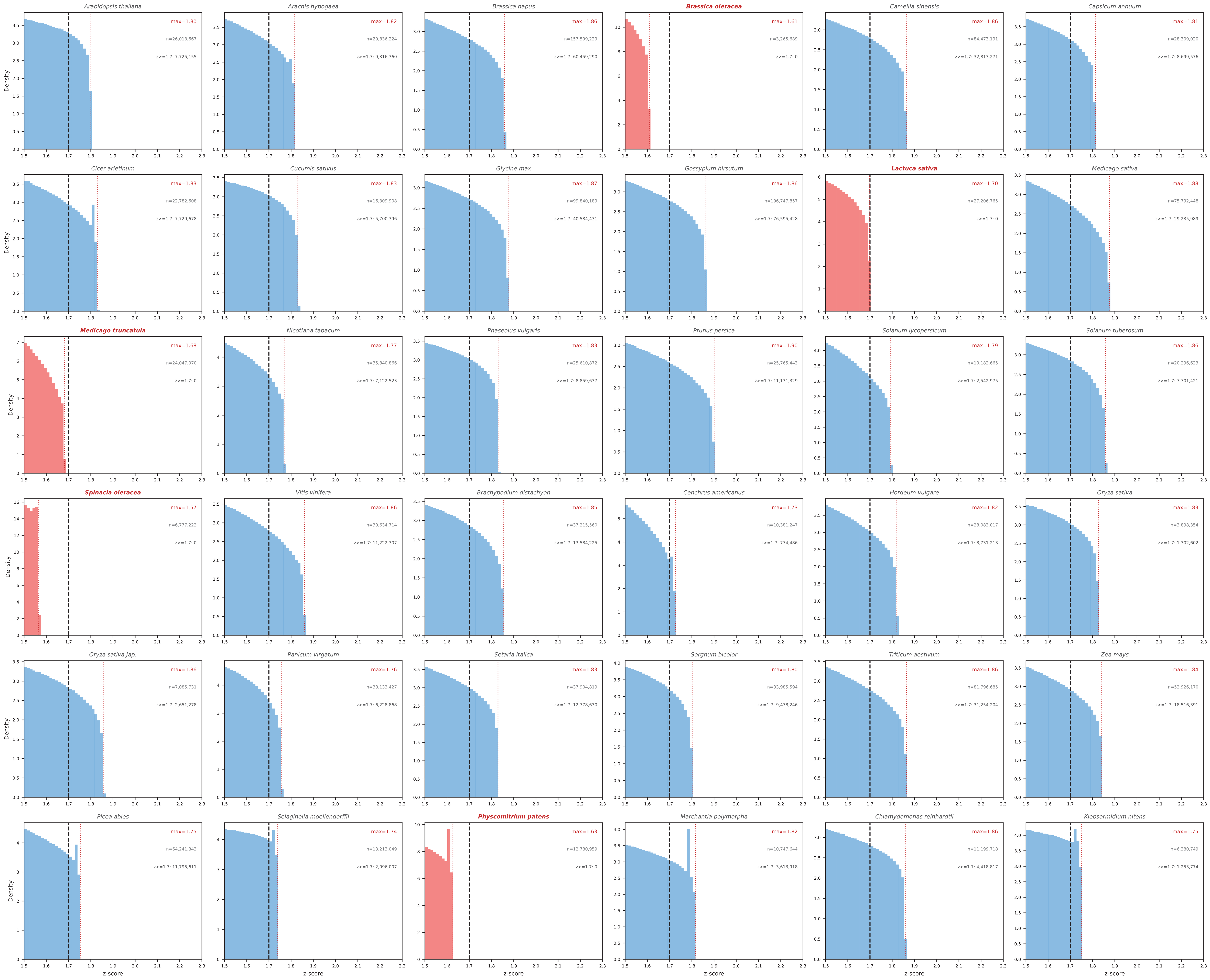


### Figure S8

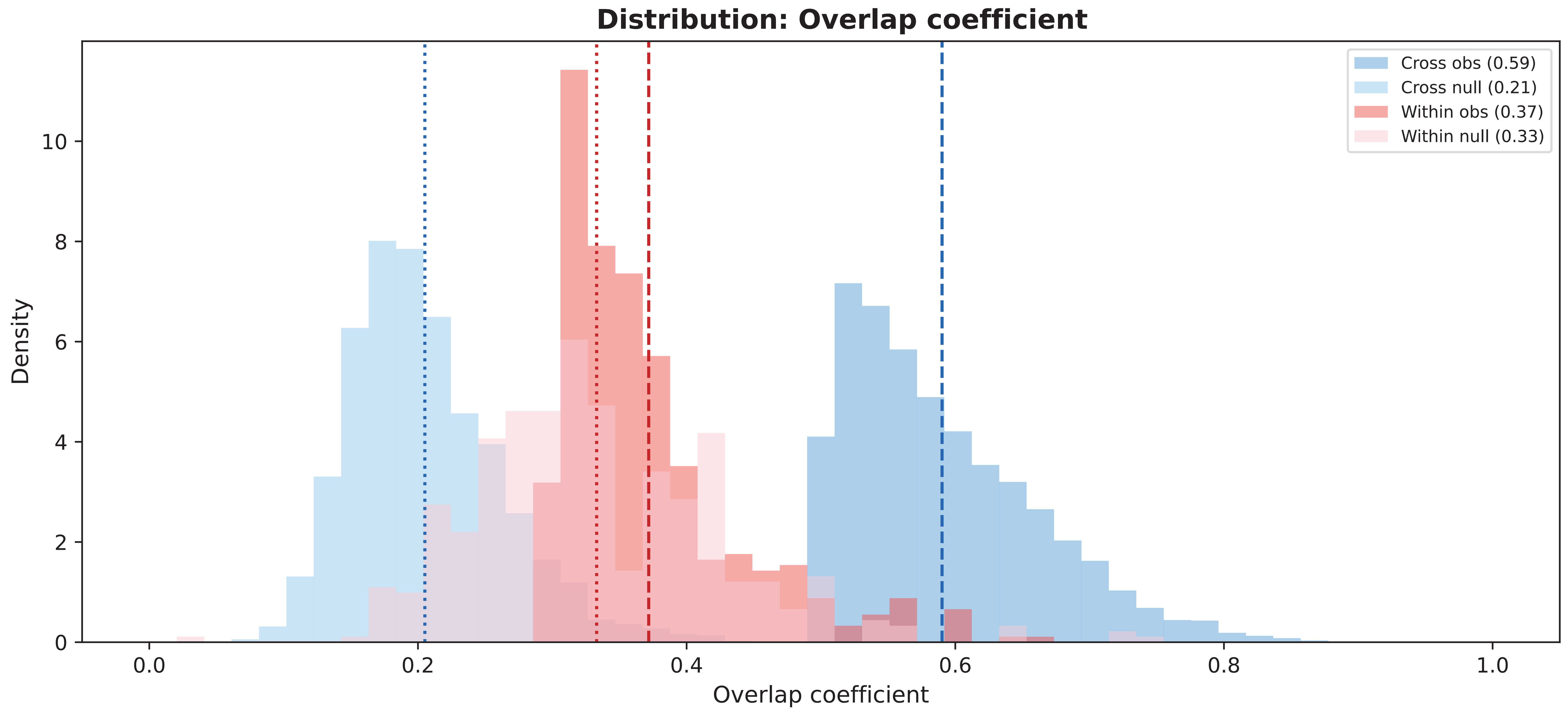
Distribution: Overlap coefficient
Cross obs (0.59)
Cross null (0.21)
Within obs (0.37)
Within null (0.33)
Density
Overlap coefficient
0
2
4
6
8
10
0.0
0.2
0.4
0.6
0.8
1.0

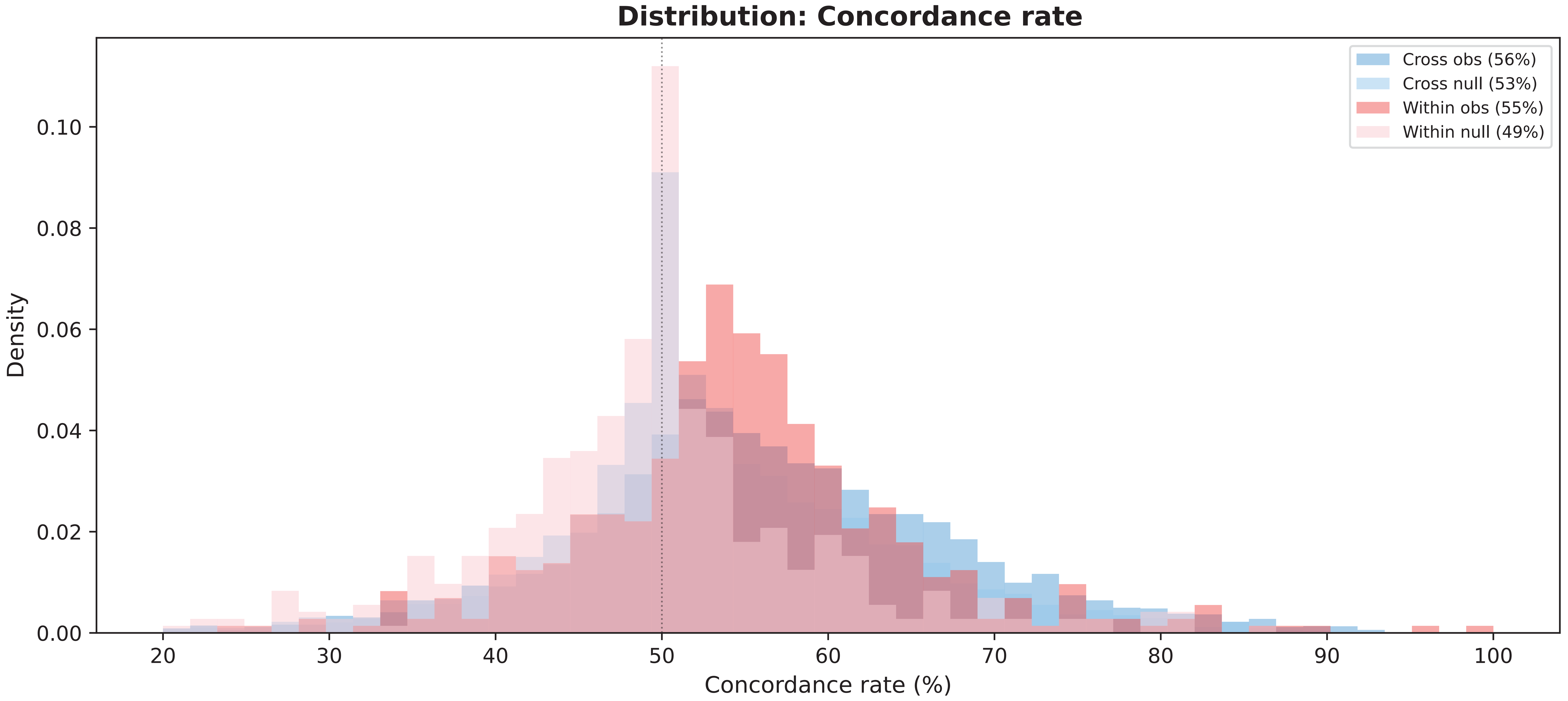
Distribution: Concordance rate
Cross obs (56%)
Cross null (53%)
Within obs (55%)
Within null (49%)
Density
Concordance rate (%)
0.00
0.02
0.04
0.06
0.08
0.10
20
30
40
50
60
70
80
90
100

Figure S9
